# Transient tertiary structure in intrinsically disordered proteins revealed by multithermal enhanced sampling

**DOI:** 10.64898/2026.01.17.700112

**Authors:** Julian O. Streit, Michele Invernizzi, Sandro Bottaro, Kamil Tamiola, Kresten Lindorff-Larsen

**Author notes:** **For correspondence:** (KLL).

## Abstract

Intrinsically disordered proteins populate heterogeneous conformational ensembles that are challenging to characterise. While all-atom molecular dynamics simulations can provide highly detailed insights into dynamic ensembles, achieving sufficient sampling remains difficult. Here, we show that On-the-fly Probability Enhanced Sampling (OPES) in the multithermal ensemble enables efficient generation of atomistic ensembles for disordered peptides and proteins ranging from 15 to 71 residues in length. Using the potential energy as a collective variable, OPES achieves multithermal sampling within a single simulation replica, without replica exchange or extensive parameter tuning. Across multiple systems, OPES yields reweighted ensembles broadly consistent with replica-exchange with solute tempering (REST2) and unbiased simulations, while accelerating convergence and enabling broader exploration of low-population conformational states. Applied to the intrinsically disordered transcriptional coactivator ACTR, OPES reveals transiently structured states in which multiple *α*-helices involved in partner binding fold cooperatively and form tertiary contacts. These rare, partially structured conformations are reversibly sampled during the simulations, consistent with extensive NMR and SAXS data, and could facilitate folding-upon-binding through conformational selection. They may also represent viable targets for drug design or for engineering disordered proteins with customised conformational landscapes. More broadly, our results establish OPES multithermal sampling as a robust and accessible approach for uncovering rare, functionally relevant conformations in intrinsically disordered proteins.

## Introduction

Understanding intrinsically disordered proteins (IDPs) and linking their conformational properties to their molecular functions remains a central challenge in molecular biophysics. Approximately one third of all protein residues in eukaryotic proteomes are in IDPs or within intrinsically disordered regions (IDRs) (***Pancsa and Tompa, 2012***), and their molecular functions underpin fundamental cellular processes ranging from transcription to signalling (***Holehouse and Kragelund, 2024***). These systems populate broad and heterogeneous conformational ensembles rather than a single, well-defined tertiary structure, making them difficult to characterise experimentally and computationally (***Borthakur et al., 2025***).

Recent methodological advances have substantially improved our ability to model these highly dynamic systems. Structure prediction methods based on deep learning, revolutionised by AlphaFold (***Jumper et al., 2021***), have begun to move beyond static structures and towards the generation of conformational ensembles that aim to explicitly capture conformational heterogeneity, including those of IDPs (***Janson et al., 2023***; ***Lewis et al., 2025***; ***Invernizzi et al., 2025***; ***Zhang et al., 2025***; ***Zhu et al., 2025***; ***Schnapka et al., 2025***; ***Novak et al., 2025***). In parallel, improvements in all-atom explicit solvent force fields have markedly enhanced the accuracy with which molecular dynamics (MD) simulations reproduce secondary-structure propensities and global compaction levels in disordered proteins (***Best et al., 2014***; ***Robustelli et al., 2018***; ***Phan et al., 2025***). A key advantage of MD over most current generative AI methods (***Lewis et al., 2025***; ***Invernizzi et al., 2025***) is its ability to probe the response of disordered states to environmental perturbations and to model multiple biomolecules simultaneously, thereby mimicking aspects of the native cellular context (***Galvanetto et al., 2023***; ***Streit et al., 2024***). However, achieving statistically robust sampling of disordered states with unbiased MD remains computationally demanding (***Lindorff-Larsen et al., 2012b; Borthakur et al., 2025***; ***Koneru et al., 2025***; ***Muhammedkutty et al., 2025***). This is particularly the case for long sequences or conditions requiring large solvation boxes, where the conformational space grows rapidly and longer simulation times are required for adequate sampling.

To overcome these challenges, all-atom implicit solvent (***Vitalis and Pappu, 2009***; ***Bottaro et al., 2013***; ***Greener, 2024***) and coarse-grained (CG) models (***Klein et al., 2021***; ***Regy et al., 2021***; ***Joseph et al., 2021***; ***Tesei and Lindorff-Larsen, 2023***; ***Cao et al., 2024***; ***Thomasen et al., 2024***; ***Wang et al., 2025***; ***Jussupow et al., 2025***) have been developed to balance accuracy, resolution and sampling efficiency. In particular, single–bead-per-residue models enable proteome-scale characterisation of IDPs at tractable computational cost (***Tesei et al., 2024***). However, their reduced resolution limits the quantitative description of local structural features, transient secondary and tertiary structure, and interactions with binding partners, ions, or small molecules, which are all of increased interest (***Borgia et al., 2018***; ***Lohr et al., 2022***; ***Kjaer et al., 2025***).

Enhanced sampling methods in all-atom explicit-solvent simulations enable efficient exploration of the high-dimensional free-energy landscapes of IDPs. Conventional temperature replica-exchange MD (TREMD) is often impractical for full-length IDPs because it requires a prohibitively large number of replicas to ensure sufficient energy overlap, although it has been successfully applied to smaller IDP fragments that can be assembled using hierarchical chain-growth strategies (***Pietrek et al., 2019***). These limitations have motivated the more widespread adoption of replica-exchange with solute tempering (REST2) (***Wang et al., 2011***; ***Bussi, 2014***; ***Shrestha et al., 2019***; ***Zhu et al., 2022***; ***Koneru et al., 2025***). Despite notable successes, including applications to IDP–small molecule interactions (***Zhu et al., 2022***; ***Zhu and Robustelli, 2025***), REST2 and related replica-based approaches face persistent challenges such as poor replica mixing, artificial compaction of disordered ensembles at high effective temperatures (***Zhang et al., 2023***), complicated implementation methods, requiring careful consideration of the temperature spacing between replicas, and substantial computational cost for large systems. Recent refinements, including REST3 (***Zhang et al., 2023***) or additional modest heating of the temperature bath (***Appadurai et al., 2021***), can partially mitigate issues related to poor replica mixing but still require careful, system-specific considerations of simulation parameters. Other enhanced sampling strategies, such as metadynamics (***Laio and Parrinello, 2002***), also introduce technical complexity, which requires careful system-specific consideration of collective variables and long simulation times to reach convergence (***Heller et al., 2020***; ***Pesce and Lindorff-Larsen, 2023***).

In this work, we introduce On-the-fly Probability Enhanced Sampling (OPES) (***Invernizzi and Parrinello, 2020***; ***Invernizzi et al., 2020***) as a practical and robust approach for generating conformational ensembles of IDPs. By using OPES with the potential energy as a collective variable, a single simulation replica can efficiently sample a multithermal ensemble (OPES multiT) with a converged bias potential. This results in ensembles that can be reweighted at multiple temperatures and avoids the need for replica exchange, alternative temperature scaling protocols, and the challenges associated with estimating accurate weights in simulated tempering simulations (***Stratmann et al., 2025***). Our results show that OPES produces reweighted ensembles consistent with REST2 and unbiased MD, while improving convergence, sampling efficiency, and exploration of conformational space across a range of systems. For the longest protein studied here, the activator for thyroid hormone and retinoid receptors (ACTR, 71 residues), OPES simulations reveal transiently formed states with tertiary structure not captured by REST2 or unbiased simulations. These results, obtained without using a structure-based reaction coordinate, highlight the importance of thorough conformational sampling and demonstrate the potential of OPES to offer new structural insights into the ensemble–function relationship of IDPs. All data and analysis code are freely available to the community (see Code/Data Availability).

## Results

### Overview of the approach and model systems

In this study, we simulated four different systems of increasing size and complexity in explicit solvent using a multithermal ensembles sampled with OPES (Figure 1). The simplest system is the short helical peptide consisting of 15 amino acids, Ace-(AAQAA)_3_-NMe (simply referred to as (AAQAA)_3_ here). It has been widely adopted by the force-field community to benchmark the ability of force fields to capture the temperature-dependent helix-coil balance of peptides (***Best and Hummer, 2009***; ***Lindorff-Larsen et al., 2012a***; ***Best et al., 2014***; ***Huang and MacKerell, 2014***; ***Robustelli et al., 2018***; ***Phan et al., 2025***). We also simulated two IDPs of 66 and 71 residues, namely the first exon of human huntingtin with a polyglutamine repeat length of 16 (HTTex1 16Q) and human ACTR, respectively—as well as a fragment of ACTR. The system sizes ranged from 29,730 ((AAQAA)_3_) to 174,602 atoms (HTTex1 16Q) and the detailed setups are described in the Methods section (Table 1). REST2 simulations were also performed for all systems to assess whether these methods yield comparable conformational ensembles, with a total aggregate simulation time equivalent to or longer than the OPES simulations.

**Table 1.**
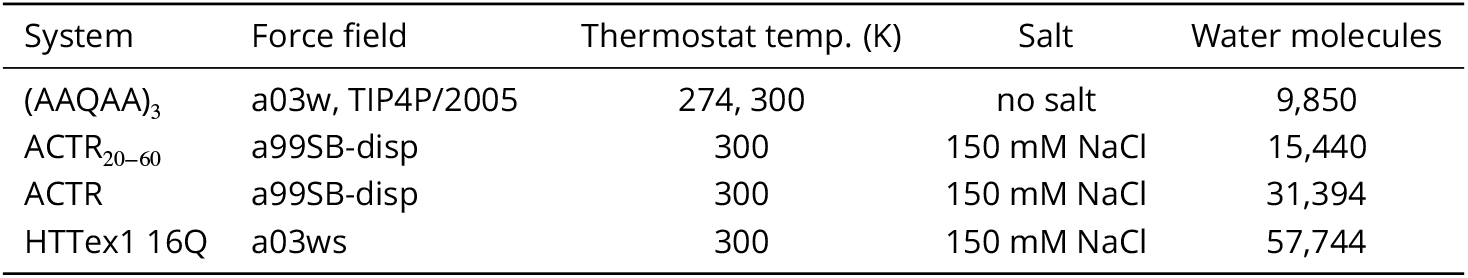
Overview of simulation systems, force fields, thermostat temperatures, salt concentrations and number of water molecules in the simulation boxes. Force-field references: ff03w (***Best and Mittal, 2010***), TIP4P/2005 (***Abascal and Vega, 2005***), a99sb-disp (***Robustelli et al., 2018***), ff03ws (***Best et al., 2014***). HTTex1 simulations used the modified Lennard Jones for Na^+^ and Cl^−^ from (***Luo and Roux, 2010***).

**Figure 1.**
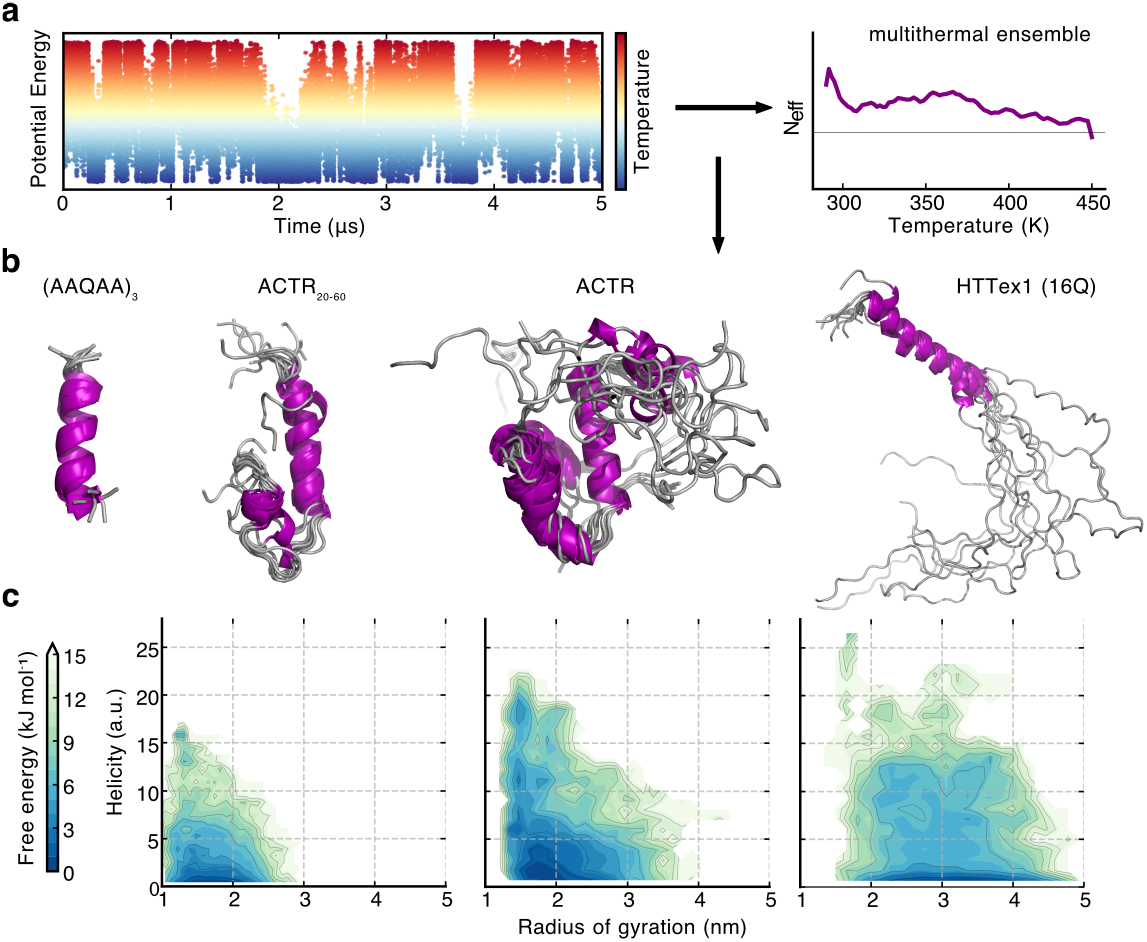
Sampling disordered peptides and proteins with OPES multithermal simulations. **a.** Representative multithermal molecular dynamics simulation showing fluctuations in potential energy (left) as the system explores a broad temperature range (colour scale). The right panel shows the relative effective sample size, *N*_eff_, sampled across temperatures, demonstrating relatively uniform ensemble coverage. The black horizontal line represents the expected sample size in an equivalent temperature replica exchange setup (1 / M, where M is the number of temperatures). **b**. Structures of the most helical conformations sampled during multithermal simulations for four systems of increasing complexity: the helical control peptide (AAQAA)_3_, the ACTR_20−60_ fragment, full-length ACTR, and HTTex1 with 16 glutamine repeats (16Q). **c**. Free-energy landscapes plotted as a function of helicity (see Methods) and radius of gyration for the ACTR_20−60_ fragment, full-length ACTR and HTTex1 16Q at 300 K.

(AAQAA)_3_ was chosen due to its smaller size, thus enabling quantitative comparisons with unbiased and REST2 simulations performed under equivalent conditions. HTTex1 and ACTR were selected because they represent larger, more realistic IDPs, associated with significant challenges for sampling: HTTex1 is an aggregation-prone protein with helical structure in its N-terminal N17 region, which can further propagate into the polyglutamine (polyQ) tract to form long helices stabilised by both backbone and side-chain hydrogen bonds (***Elena-Real et al., 2023***; ***Mohanty et al., 2025***), which is a general feature of polyQ sequences (***Escobedo et al., 2019***). It is also relatively extended due to its C-terminal proline-rich region, thus making the convergence of both secondary structure and extent of compaction challenging. The longest IDP studied in this work is human ACTR (71 residues), which acts as a transcriptional coactivator by binding to the nuclear coactivator binding domain (NCBD) of CREB binding protein (CBP). Upon binding, ACTR and NCBD fold into a stable structure each contributing three *α*-helices (***Demarest et al., 2002***). In the absence of NCBD, ACTR remains disordered but samples residual helical structure in helix 1 (residues 27– 41) (***Kjaergaard et al., 2010a***; ***Iešmantavičius et al., 2013***). Additionally, we considered whether a smaller fragment of an IDP may be used as a suitable alternative to systematically explore regions for structural propensity at reduced computational cost. To this end, we performed simulations of a fragment of ACTR consisting of residues 20–60 (ACTR_20−60_), which includes the region of helix 1 folded in the ACTR-NCBD complex.

For each system, we initially performed ten independent, short trial simulations to build a bias potential with the potential energy as the collective variable (CV, ‘bias convergence simulations’, see Methods). The OPES bias potentials generally converged within 100 and 600 ns for the smallest and largest systems studied, respectively (Figure S1). Converged bias potentials were subsequently used as static biases in multiple production simulation replicas to increase throughput (see Methods). Unlike replica exchange-based methods, the OPES production replicas are non-exchanging and use the same bias potential. Consequently, the systems diffuse through a predefined temperature range as their potential energies fluctuate during the simulations, resulting in a multithermal ensemble with approximately uniform effective sample sizes across all temperatures (Figure 1a). This enabled sampling of heterogeneous conformational ensembles, ranging from highly compact and helical states (Figure 1b) to more extended conformations lacking secondary structure (Figure 1c).

### Helix-coil transitions of the (AAQAA)_3_ peptide

We compared the ensembles and convergence of the (AAQAA)_3_ peptide generated by REST2 and OPES multiT simulations at 300 K. For both methods, we observed roughly uniform sampling of temperature space (Figures S2, S3, S4). Time series of the radius of gyration (Rg) and helicity (Figure 2a) show that both methods effectively sample the conformational space over 15 *μ*s, with numerous reversible transitions between disordered and fully helical states. Probability distributions of R_*g*_ (Figure 2b) indicate a largely overlapping ensemble between REST2 and OPES, with mean values close to 1 nm and within error of each other, suggesting that both methods yield similar global conformational properties. Residue-wise helicity profiles (Figure 2c) further confirm this agreement, capturing the characteristic helical propensity of the central residues, consistent across both methods. To quantify the efficiency of sampling, we considered multiple metrics. An autocorrelation analysis (Figure 2d) reveals that OPES accelerates the decorrelation of both R_*g*_ and helicity by a factor of 2.0–2.5 relative to REST2, indicating enhanced sampling efficiency. A similar acceleration factor is obtained when comparing the error bars from a blocking analysis of both ensembles: from the ratio of the error bars, OPES simulations appear to sample conformational space 2–3× more efficiently (Figure 2d). Time-cumulative averages of R_*g*_ and the total helical fraction (Figure 2e) also demonstrate more rapid convergence for the latter observable in OPES relative to REST2, highlighting the ability of OPES multiT to reach equilibrium faster while reproducing the same structural ensemble. Collectively, these results suggest that OPES multiT offers improved sampling efficiency for peptide folding simulations.

**Figure 2.**
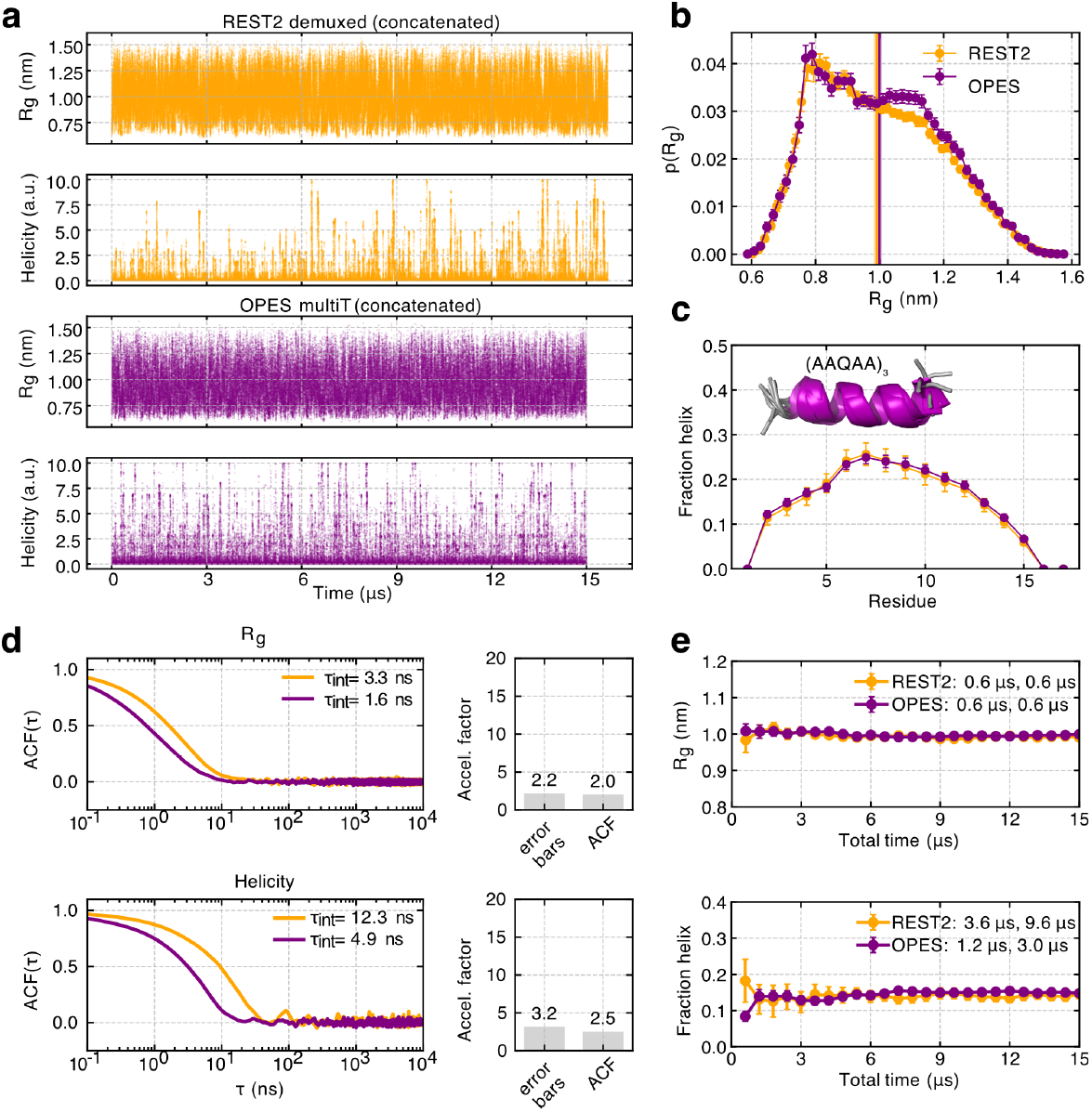
Sampling efficiency and convergence rate of OPES multiT and REST2 simulations of the helical (AAQAA)_3_ peptide. **a.** Fluctuations of the radius of gyration (R_*g*_) and peptide helicity sampled with REST2 and OPES multiT simulations. **b**. Probability distributions and ensemble-averaged values (vertical lines) of the R_*g*_. Errors represent the standard error of the mean calculated with a blocking analysis (see Methods). **c**. Mean and standard errors of per-residue helical fractions. **d**. Normalised autocorrelation function (ACF) of the R_*g*_ (top) and helicity (bottom) calculated from the concatenated REST2 and OPES multiT timeseries sampled every 40 ps. The correlation times were estimated using the integral of the ACF (*τ*_*int*_). The right panels show estimates of an acceleration factor (OPES / REST2) estimated from the error bars of ensemble-averaged values and correlation times (ACF). **e**. Time-cumulative averages are shown together with the extracted stabilisation times, which mark the earliest points at which the running averages and their uncertainties, respectively, converge to stable values within predefined tolerances (see Methods).

We assessed the reproducibility of OPES multiT simulations using independent bias potentials at a reference temperature of 300 K. Ensembles generated from three replicas with independent biases were highly consistent with those obtained using a shared bias potential, with only minor, localised differences observed in the R_*g*_ probability distribution between 1.1 and 1.2 nm (Figure S2e). A similarly small discrepancy was also observed when comparing REST2 and OPES distributions (Figure 2b), suggesting that these deviations arise from finite sampling and the possible under-estimation of errors obtained with blocking (see Methods and Figure S1c) rather than systematic differences between methods. Simulations performed at 274 K, when reweighted to 300 K, also yielded distributions broadly consistent with those obtained directly at 300 K (Figure S2f). Ensembles generated using independent biases at 274 K were likewise consistent with those obtained using a shared bias potential (Figure S3e). Taken together, these results indicate that the enhanced sampling simulations of (AAQAA)_3_ are converged with respect to the average R_*g*_, overall shape of the R_*g*_ distribution and secondary structure content, with only small, localised differences in the R_*g*_ distributions that are unlikely to affect the overall conclusions regarding sampling efficiency.

To quantify the acceleration of sampling achieved by OPES compared to unbiased MD, we performed 15 *μ*s unbiased simulations of (AAQAA)_3_ at 274 and 300 K. The time series of R_*g*_ and helicity at 300 K show multiple helix–coil transitions, indicative of good sampling even in unbiased simulations (Figure S5a,f). The R_*g*_ probability distributions, average R_*g*_, and per-residue helical propensities are all consistent within error (Figure S5b–c). Based on the error bars of ensemble-averaged properties, OPES simulations accelerate sampling by roughly 3–6×, depending on the property analysed (Figure S5d). Autocorrelation analyses suggest even higher acceleration factors of 13– 16×, indicating that diffusion through conformational space is enhanced by an order of magnitude (Figure S5d). Time-cumulative averages further reveal that helical propensities stabilise earlier in OPES simulations than in unbiased MD, by approximately 3–5× (Figure S5e).

Results at 274 K show that ensembles generated by unbiased MD and OPES are broadly consistent in terms of R_*g*_ and per-residue helical propensities. OPES simulations also exhibit a clear acceleration relative to unbiased MD, with factors of approximately 11–18× based on error bars, 35–79× based on autocorrelation analyses, and 3× based on time-cumulative helical propensities (Figure S6). Overall, these findings validate that OPES multiT produces reproducible ensembles that reflect the force field while offering improved sampling efficiency compared to both unbiased MD and REST2 simulations of (AAQAA)_3_.

### Efficient sampling of an extended IDP with helical structure

Next, we performed REST2 and OPES multiT simulations of a larger IDP system, HTTex1 16Q (Figure 3a). For the OPES simulations, we employed ten non-exchanging replicas of 2 *μ*s each, for a total aggregated sampling time of 20 *μ*s. All replicas showed efficient diffusion in temperature space and relatively uniform sampling across the 288–450 K range (Figure S7a). Consistent with this, the R_*g*_ and helicity fluctuated throughout the simulations, indicating adequate exploration of conformational space. The N-terminal region, which is known to exhibit a high helical propensity (***Elena-Real et al., 2023***), underwent multiple helix–coil transitions with helices of varying lengths (Figure S7b), in agreement with previous experimental and computational studies (***Elena-Real et al., 2023***; ***Mohanty et al., 2025***).

**Figure 3.**
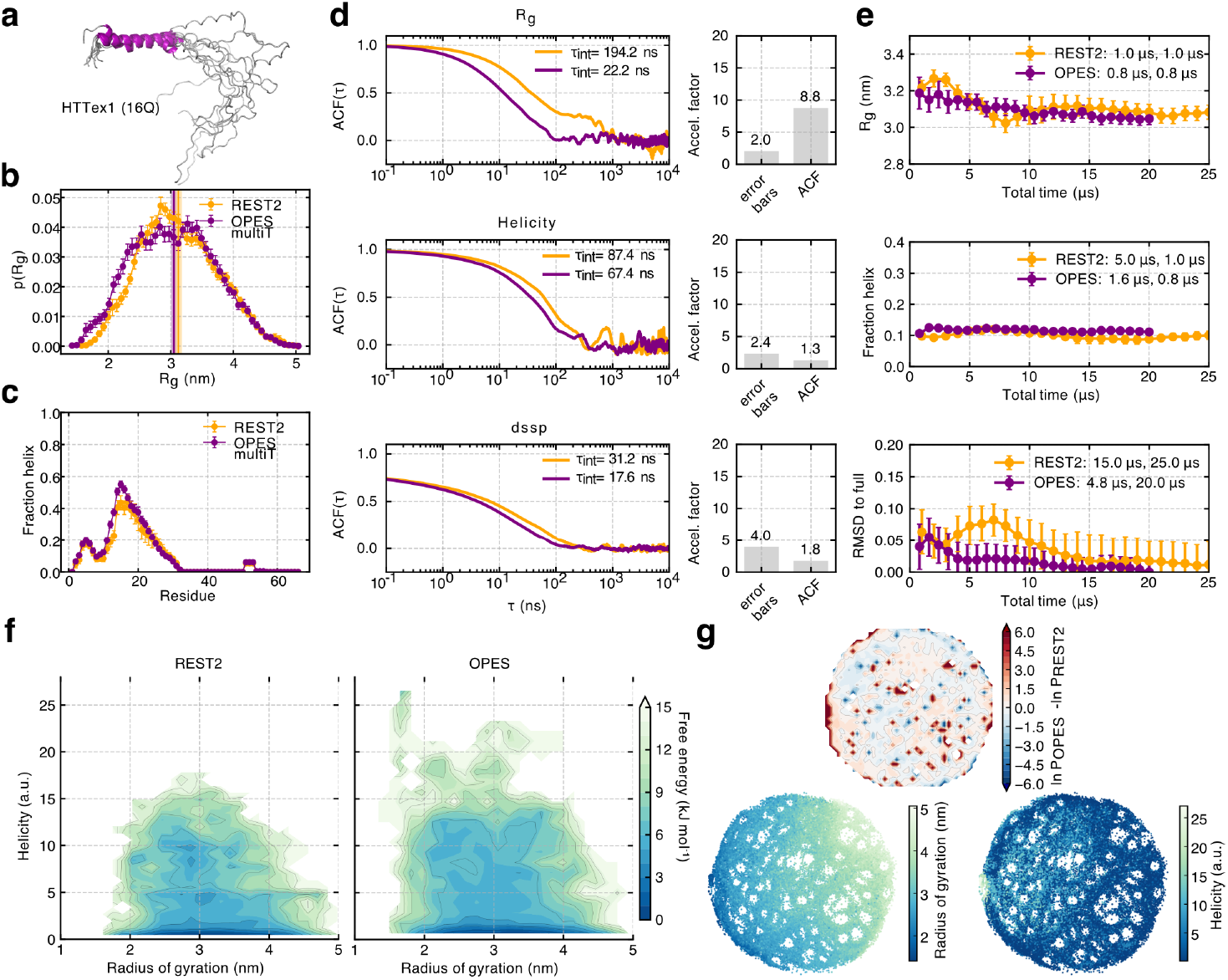
OPES multiT and REST2 simulations of HTTex1 16Q. **a.** Exemplar structures of the most helical state sampled with OPES multiT. **b**. Probability distributions and ensemble-averaged values (vertical lines) of the R_*g*_. Errors represent the standard error of the mean. **c**. Mean and standard errors of per-residue helical fractions. **d**. Normalised autocorrelation function (ACF) of the R_*g*_ (top), helicity (bottom) and per-residue secondary structure (dssp, see Methods) calculated from the concatenated REST2 and OPES multiT timeseries sampled every 40 ps. The correlation times were estimated using the integral of the ACF (*τ*_*int*_). The right panels show estimates of an acceleration factor (OPES / REST2) estimated from the error bars of ensemble-averaged values and correlation times (ACF). **e**. Time-cumulative averages of the R_*g*_, total fraction helix and per-residue helical fractions (measuring the RMSD to the profile derived from the full ensemble) are shown together with the extracted stabilisation times, which mark the earliest points at which the running averages and their uncertainties, respectively, converge to stable values within predefined tolerances. **f**. Free-energy landscapes as a function of helicity and R_*g*_ at 293 K. **g**. 2D ELViM projections of a combined REST2 + OPES ensemble at 293 K (see Methods). Top: difference in log probabilities across the entire landscape. Bottom: projections coloured by R_*g*_ and helicity.

REST2 simulations similarly demonstrated efficient temperature-space diffusion, with comparable numbers of round trips across most replicas. One exception was replica 10, which became trapped for a substantial portion of the simulation at high effective temperature in a highly compact conformational state (Figure S8a–b). Despite this, the majority of replicas sampled helix–coil transitions and a broad range of conformations (Figure S8c–d).

Figure 3 summarises the comparative conformational ensembles and sampling efficiency obtained for HTTex1 16Q using REST2 and OPES multiT simulations. Both methods yield similar equilibrium ensembles, as evidenced by similar R_*g*_ probability distributions (Figure 3b) and comparable ensemble averages (3.05 ± 0.03 nm and 3.12 ± 0.05 nm for OPES and REST2, respectively). Per-residue helical propensities are in good agreement, with both approaches sampling helical structure in the same regions of the sequence. However, OPES simulations display approximately 10% higher helical content at the peak of the second helical region (Figure 3c), indicating slightly enhanced exploration of helical states.

In terms of sampling efficiency, OPES exhibits systematically faster dynamics. Autocorrelation function (ACF) analyses of R_*g*_, total helical content, and DSSP-based secondary structure reveal shorter integrated correlation times for OPES relative to REST2 (Figure 3d), corresponding to acceleration factors of approximately 1.3–8.8× depending on the observable. Convergence analyses further show that OPES reaches stable estimates the total helical fraction and RMSD with respect to the per-residue helicities of the full ensembles at shorter accumulated simulation times than REST2 (Figure 3e), with an improvement of roughly a factor of three for both total and per-residue helicity.

Two-dimensional (2D) free-energy surfaces projected onto R_*g*_ and helicity demonstrate that both methods sample a highly heterogeneous and largely overlapping region of conformational space (Figure 3f). For larger IDPs such as HTTex1, it is therefore important to assess not only convergence but also the extent of conformational space exploration. To this end, we performed an additional data-driven dimensionality-reduction analysis using ELViM (***Viegas et al., 2024***), which projects conformational ensembles onto a two-dimensional grid based on internal C*α* distances as a similarity measure. These projections separate states primarily according to R_*g*_ and appear to be less sensitive to secondary structure (Figure 3g).

When projected onto the same ELViM space, OPES and REST2 display comparable effective coverage and entropy, indicating similar exploration of the dominant conformational basins (Table S3). In contrast, projections into a shared R_*g*_–helicity space reveal broader sampling by OPES. In this representation, OPES exhibits approximately 30% higher effective coverage and 5% higher entropy, and samples nearly all bins accessed by REST2, whereas REST2 explores approximately 35% fewer bins overall (Table S4). This suggests that OPES more effectively accesses low-population helical states that are visited less frequently in REST2 simulations. Overall, these results demonstrate that while REST2 and OPES generate comparable equilibrium ensembles for HTTex1 16Q, OPES achieves broader exploration of conformational states with varying compactness and secondary structure, alongside moderately faster convergence.

### OPES enables improved conformational space exploration

As a second challenging target for all-atom MD simulations, we studied the 71-residue human ACTR, an intrinsically disordered protein (IDP) that folds upon binding (***Demarest et al., 2002***) and displays residual helicity in the apo state (***Kjaergaard et al., 2010a***; ***Iešmantavičius et al., 2013***). Likely owing to functional constraints associated with binding, ACTR is more hydrophobic than HTTex1 16Q (GRAVY score (***Kyte and Doolittle, 1982***; ***Gasteiger et al., 2005***) −0.672 vs. −1.473). Despite lacking aromatic residues, this increased hydrophobicity arises from a high aliphatic content, reflected in an aliphatic index (***Ikai, 1980***) of 102 compared to 40 for HTTex1 16Q. Most aliphatic residues (predominantly leucine) are located at the CBP-binding interface, conferring amphipathic character to the three helical regions involved in binding (residues 27–41, 46–54, and 56–63). Consequently, ACTR is expected to exhibit a slightly more rugged free-energy landscape than the low-complexity HTTex1 sequence.

We first simulated a 40-residue fragment (ACTR_20−60_) encompassing the primary helix that exhibits residual structure in the apo state (helix 1, residues 27–41). OPES multiT simulations were performed using five replicas of 5 *μ*s each, with two independent bias potentials to assess reproducibility. In both cases, replicas diffused efficiently through temperature space, exhibiting persistent fluctuations in R_*g*_, helicity and near-uniform temperature sampling (Figure S9a–d). Numerous helix–coil transitions were observed throughout the sequence, indicating robust secondary-structure sampling (Figure S9e). The two OPES biases yielded broadly consistent ensembles, with similar R_*g*_ distributions (mean R_*g*_ = 1.68 ± 0.01 and 1.64 ± 0.01 nm), per-residue helical propensities, and temperature-dependent trends (Figure S9f). As observed for (AAQAA)_3_, some small, localised differences exceeding the combined uncertainties are present, for example in per-residue helicities and in limited regions of the R_*g*_ distributions.

REST2 simulations of ACTR_20−60_ also showed good temperature diffusion and sampling of a broad range of R_*g*_ values and helix–coil transitions (Figure S10). As observed for HTTex1, OPES and REST2 produced similar ensembles in terms of R_*g*_ and per-residue helicity, although OPES yielded a slightly higher overall helical content (Figure S11a-c). Autocorrelation analyses indicate faster sampling with OPES, with acceleration factors of 4.4–6.5× and earlier stabilisation of per-residue helicity by approximately a factor of two (Figure S11d–e). However, acceleration estimates based on final error bars are close to unity. Examination of free-energy landscapes and ELViM projections appears to partially reconcile this discrepancy: while both methods exhibit similar effective coverage and normalised entropy in ELViM space (Table S5), OPES samples ~40% more area in R_*g*_–helicity space (Table S6). This reflects additional free-energy minima corresponding to highly helical states (helicity > 12.5) sampled only by OPES (Figure S11f). Thus, OPES enhances sampling speed and access to low-population helical states while yielding comparable ensemble averages and uncertainties.

Encouraged by these results, we simulated the full-length 71-residue ACTR. Despite its larger size, OPES multiT achieved efficient temperature-space exploration in 5 replicas of 5 *μ*s, with frequent helix-coil transitions and relatively uniform temperature sampling (Figure S12a–c). This produced smooth temperature-dependent trends of the average R_*g*_ and helicity, and per-residue helical profiles consistent with ACTR_20−60_, except for helix 3, which lacks its full sequence in the fragment (Figure S12d). In contrast, REST2 simulations exhibited substantially reduced temperature diffusion, with an order-of-magnitude decrease in replica round trips relative to HTTex1 and ACTR_20−60_ (Figure S13), consistent with prior observations for larger IDPs where artificially collapsed replicas get trapped at high effective temperatures (***Zhang et al., 2023***). Nevertheless, final R_*g*_ distributions and per-residue helicities from REST2 and OPES agree reasonably well (Figure 4b–c).

**Figure 4.**
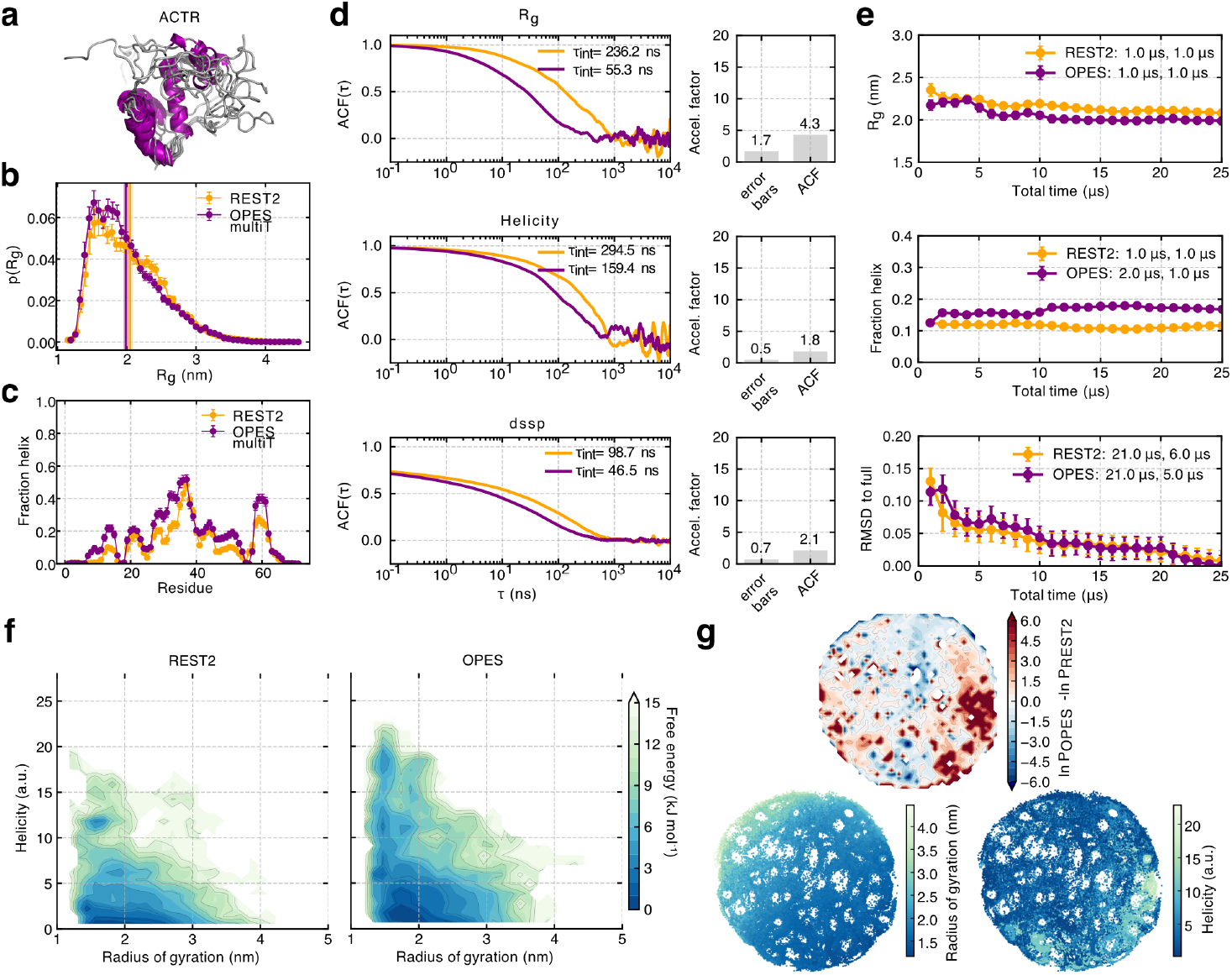
OPES multiT and REST2 simulations of ACTR. **a.**Exemplar structures of the most helical state sampled with OPES multiT. **b**. Probability distributions and ensemble-averaged values (vertical lines) of the R_*g*_. Errors represent the standard error of the mean. **c**. Mean and standard errors of per-residue helical fractions. **d**. Normalised autocorrelation function (ACF) of the R_*g*_ (top), helicity (bottom) and per-residue secondary structure (dssp, see Methods) calculated from the concatenated REST2 and OPES multiT timeseries sampled every 40 ps. The correlation times were estimated using the integral of the ACF (*τ*_*int*_). The right panels show estimates of an acceleration factor (OPES / REST2) estimated from the error bars of ensemble-averaged values and correlation times (ACF). **e**. Time-cumulative averages of the R_*g*_, total fraction helix and per-residue helical fractions (measuring the RMSD to the profile derived from the full ensemble) are shown together with the extracted stabilisation times, which mark the earliest points at which the running averages and their uncertainties, respectively, converge to stable values within predefined tolerances. **f**. Free-energy landscapes as a function of helicity and R_*g*_ at 300 K. **g**. 2D ELViM projections of a combined REST2 + OPES ensemble at 300 K (see Methods). Top: difference in log probabilities across the entire landscape. Bottom: projections coloured by R_*g*_ and helicity.

Autocorrelation analyses for full-length ACTR again show modestly faster sampling with OPES (1.8–4.3×), whereas error bar-based acceleration factors are closer to unity and ensemble averages stabilise at similar rates (Figure 4d–e). Notably, projections onto R_*g*_-helicity and ELViM space reveal broader conformational exploration by OPES, including additional free-energy minima corresponding to highly helical states (helicity > 14.0) not accessed by REST2 (Figure 4f–g). OPES exhibits higher effective coverage (30–60%), normalised entropy, and more complete overlap with REST2-sampled regions (Tables S7, S8).

Finally, we compared OPES multiT simulations with a previously published 30 *μ*s unbiased MD simulation using the same force field (***Robustelli et al., 2018***). Differences in simulation setup preclude a strict one-to-one comparison (***Piana et al., 2012***), but both error-bar and autocorrelation analyses indicate OPES acceleration factors of 3.1–5.9×, depending on the collective variable (Figure S14d–f), consistent with time-cumulative helicity estimates (Figure S14e). As with REST2, unbiased simulations do not sample the highly helical states observed with OPES (Figure S14g). Overall, these results demonstrate that OPES multiT enables robust and efficient sampling of realistic IDPs, achieving comparable or improved convergence relative to REST2 and unbiased MD while accessing a broader conformational landscape, particularly low-population compact helical states.

### An integrative structural ensemble of ACTR

Given the established role of ACTR folding-upon-binding to the NCBD of CBP in mediating its function (Figure 5a), we next asked whether the apo ensemble of ACTR samples conformations resembling the bound state. To address this, we leveraged the extensive experimental information available for ACTR. NMR chemical shifts (CS) and residual dipolar couplings (RDCs) primarily report on local structure and secondary-structure propensities (***Iešmantavičius et al., 2013***), whereas paramagnetic relaxation enhancement (PRE) NMR and small-angle X-ray scattering (SAXS) probe long-range tertiary contacts and global dimensions (***Kjaergaard et al., 2010a***; ***Iešmantavičius et al., 2013***).

**Figure 5.**
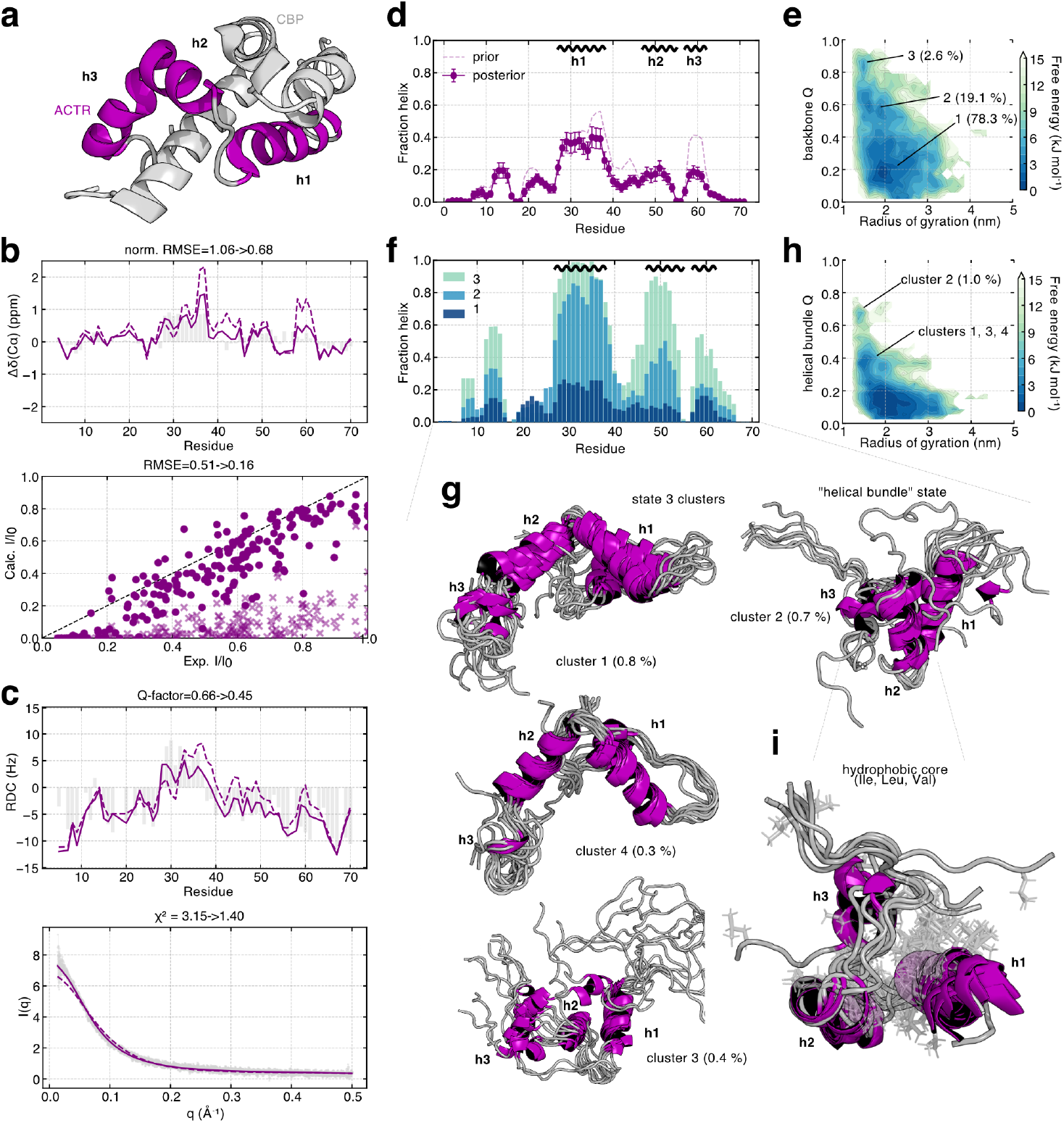
Transient formation of tertiary structure in ACTR and ensemble reweighting with experimental data. **a.** Structure of ACTR bound to its binding partner, CBP (PDB 1KBH). The three helices of ACTR involved in the interaction are annotated. **b**. Agreement with active data used for ensemble reweighting (top: secondary C*α* chemical shifts, bottom: PRE intensity ratios) before and after reweighting. **c**. Agreement with passive data excluded from ensemble reweighting (top: RDCs, bottom: SAXS) before and after reweighting. **d**. Ensemble-averaged helical propensity across the sequence of ACTR before and after reweighting (mean ± s.e.m.). The three helices of ACTR involved in the CBP interaction are annotated at the top. **e**. Free-energy landscape as a function of backbone native contacts (Q) relative to the bound state and R_*g*_ at 310 K after reweighting. The three main low energy basins are annotated with their respective populations. **f**. Helical propensities for the three low energy basins annotated in panel e. **g**. Structures of the four main conformational clusters present in state (basin) 3 from panel e. The top 10 structures by weight are shown (after reweighting). Structures were aligned using backbone atoms of residues 27–63. **h**. Free-energy landscape as a function of native contacts (Q) using a helical bundle reference structure (Figure S16d) and R_*g*_ at 310 K after reweighting. The locations of the main clusters on this landscape from panel g are annotated. **i**. Close-up view of the helical bundle-like state (cluster 2) with hydrophobic residues (Ile, Val, Leu) shown in a transparent stick representation.

To generate ensembles consistent with both local and global experimental observables, we applied Bayesian/maximum entropy reweighting (***Bottaro et al., 2020***) to ensembles obtained from REST2 and OPES simulations (see Methods). Reweighting using CS and SAXS data, with RDC and PRE data reserved for cross-validation, yielded comparable agreement with the restrained observables for both methods; however, the OPES-derived ensemble showed improved agreement with the left-out RDC and PRE data (Figure S15a).

We additionally performed reweighting using CS and PRE data as restraints (see Methods), reserving RDC and SAXS data for cross-validation (Figure S15b). In this case, agreement with CS data was comparable to that obtained from CS+SAXS reweighting, while agreement with PREs improved substantially, as expected (RMSE 0.16 for both ensembles; Figure S15c). Importantly, agreement with the independent RDC and SAXS datasets also improved for both ensembles, supporting the accuracy of the resulting reweighted ensembles. While the OPES ensemble showed better agreement with the left-out RDCs (Q-factor 0.45 vs. 0.55), the REST2 ensemble achieved slightly better agreement with SAXS data (reduced *χ*^2^ 1.19 vs. 1.40). Notably, however, SAXS agreement is similar to that obtained from CS+SAXS reweighting (reduced *χ*^2^ 1.18 vs. 1.25), indicating that multiple conformational ensembles can explain the SAXS data, whereas the PRE data impose more restrictive constraints.

Consistent with this interpretation, the average R_*g*_ of the OPES ensemble increases from 1.90 ± 0.04 nm prior to reweighting (at 310 K) to similar values after CS+SAXS and CS+PRE reweighting (2.19± 0.06 and 2.16± 0.05 nm, respectively). We therefore focus on the OPES ensemble reweighted with CS+PRE data in the following analyses, as it provides the most balanced agreement across all experimental observables (Figures 5b–c, S15d). Although these experimental data do not unequivocally distinguish between the reweighted OPES and REST2 ensembles, our results demonstrate that the additional, highly helical states uniquely sampled by OPES are consistent with all experimental measurements considered, supporting their potential relevance within the apo conformational ensemble of ACTR.

The OPES ensemble reweighted with CS+PRE data exhibits per-residue helical propensities that are similar to those of the ensemble before reweighting (prior). The highest propensity (~40%) is observed precisely in the region corresponding to helix 1 in the bound state (Figure 5d), as noted in previous experimental studies (***Kjaergaard et al., 2010a***; ***Iešmantavičius et al., 2013***). In addition, up to 20% helical structure is also detected in regions corresponding to helices 2 and 3 of the bound complex, as well as in the N-terminal segment encompassing residues ~11-17 (Figure 5d).

### Transient folding and non-native tertiary structure of ACTR

A key advantage of the MD ensemble is that it enables more detailed insights into the distribution of conformational states underlying this ensemble-averaged helical profile, which are not directly accessible from experiments. In particular, we investigated whether the apo ensemble contains conformations in which multiple helices involved in CBP binding are formed simultaneously. To this end, we used the experimental structure of ACTR bound to CBP to define a set of backbone native contacts (backbone Q) and constructed a free-energy landscape as a function of backbone Q and the R_*g*_. This analysis reveals that the apo ensemble of ACTR is composed of distinct subensembles characterised by different degrees of native-like backbone structuring (Figure 5e).

We identified three main states along the backbone Q coordinate, that we denote states 1, 2, and 3 (Figure 5e–f). Interestingly, comparison of the per-residue helical propensities across these states reveals a progressive folding of helices 1 and 2, accompanied by partial structuring of helix 3. In state 3, which accounts for approximately 2.6% of the ensemble and exhibits ~85% of native (i.e. in the complex) backbone-backbone contacts, helices 1 and 2 are both formed, with central residues adopting a helical conformation in 90–100% of this subensemble. As expected, the helices’ termini display reduced helical propensity due to fraying (Figure 5f). Helix 3 is partially folded in this state, reaching up to ~50% helicity. Inspection of inter-residue contact maps after ensemble reweighting further shows that states 1 through 3 progressively acquire tertiary structure, manifested as an increase in long-range contacts that accompanies the buildup of secondary structure (Figure S15g). This includes contacts between the N-terminal 20 residues and helices 1 and 2, consistent with PRE NMR experiments in which the spin label is attached to residue 3 (***Iešmantavičius et al., 2013***) (Figures S15d,g). Contacts are also observed between helices 1 and 2, and to a lesser extent between helices 2 and 3. The former is consistent with the long-range contact between residues 21 and 55 detected in PRE NMR experiments (Figure S15d).

When the fraction of native contacts is calculated using all contacts (All Q), rather than only backbone-backbone contacts, a free-energy minimum is observed at Q~0.6, corresponding to a state that resembles, but is not identical to, the bound conformation (Figure S15e). These free-energy landscapes are qualitatively consistent across reweighting with CS+SAXS and CS+PRE data, both revealing a bound-like basin (Figure S15e). Thus, the apo ensemble of ACTR contains a low-population (~3%) subset of structured conformations that exhibit both secondary structure and tertiary contacts reminiscent of the bound state.

We therefore further investigated whether the bound conformation of ACTR would remain stable, in terms of secondary and tertiary structure, in the absence of CBP. Starting from the experimental structure of the ACTR-CBP complex (***Demarest et al., 2002***), we performed ten unbiased MD simulations of 5 *μ*s each. Across most simulations, the relative orientations of the three helices and the overall R_*g*_ exhibited substantial fluctuations (Figures S16a-b), indicating that the precise bound conformation is not stable without CBP. This behaviour is consistent with the fact that the majority of tertiary contacts in the complex are formed between ACTR and CBP rather than within ACTR itself (***Demarest et al., 2002***). Interestingly, despite the loss of a bound-like tertiary arrangement, the secondary structure of helices 1–3 remained relatively stable in several independent simulations. In these trajectories, ACTR sampled compact conformations that were metastable on the low-microsecond timescale (Figures S16a–b). To characterise these states in more detail, we performed conformational clustering (see Methods), which revealed that the dominant clusters retain substantial helical content and exhibit tertiary contacts between helices. In some cases, these conformations resemble a collapsed helical bundle in which all three binding helices remain folded and pack together (e.g., cluster 2, Figures S16c-d). These simulations therefore suggest that while the secondary structure characteristic of the bound state is metastable in the absence of CBP, the specific bound-state three-dimensional arrangement is not. Instead, the helices can collapse into alternative compact conformations, forming tertiary contacts that give rise to bound-like but non-native structural states.

Using the top six cluster centroids (denoted C1–C6) obtained from the analysis of the unbiased MD simulations initiated from the bound state, we defined additional sets of “native contacts” and visualised free-energy landscapes as a function of the corresponding Q variables and R_*g*_. This analysis was used to assess whether similar structural states were sampled in the OPES simulations. In the OPES ensemble, we observed distinct free-energy basins at Q ~0.6-0.65 for centroids C2 and C4, but not for the other clusters, indicating that structures similar to C2 and C4 (which merely differ by the relative orientation of the third helix) are accessed by OPES. In contrast, these states were not sampled in REST2 or in unbiased simulations (***Robustelli et al., 2018***) (Figure S17a). Further inspection of the OPES time series for these Q variables shows that these compact, helical conformations—resembling C2, C4, and the bound state—are visited multiple times in a reversible manner in three out of the five OPES simulations (Figure S17b, Supplementary movies S1-S3). This behavior supports the interpretation that these conformations correspond to bona fide metastable structural states.

Finally, we visualised the structural states sampled in the OPES simulations after ensemble reweighting by clustering state 3. Four dominant clusters capture the heterogeneity of this state, in which helices 1 and 2 are largely folded and resemble the bound conformation, while differing in their relative inter-helical orientations (Figure 5g). Notably, the second cluster closely resembles the helical bundle observed in simulations initiated from the bound state, including formation of a well-defined hydrophobic core (Figure 5g–i), suggesting that it represents a folding intermediate of the fully formed bundle (C2). Indeed, the free-energy landscape projected onto C2 Q and R_*g*_ reveals a local minimum near Q~0.7 (Figure 5h). Folding of helices 1–3 is cooperative, with joint formation occurring more frequently than expected from independent events (Figure S18), indicating that these states correspond to genuine local free-energy minima with tertiary structure. Helices 1 and 2 exhibit the strongest co-folding propensity and cooperativity, in qualitative agreement with experimental observations showing stronger chemical shift perturbations (CSPs) of helix 2, compared to other helical regions, upon mutation of helix 1 (***Iešmantavičius et al., 2013***). This cooperative behaviour is observed both before and after ensemble reweighting with experimental data, is reproduced in OPES simulations of the shorter ACTR_20−60_ fragment, and is reduced at elevated temperature (440 K, Figure S18), collectively supporting the conclusion that ACTR can transiently adopt folded, helix-bundle–like conformations at physiological temperature.

In summary, the broader conformational exploration of OPES multiT simulations of ACTR enabled the detection of rare folding events characterised by hydrophobic interactions between amphipathic helices. The structural resemblance of these states to the bound conformation, their agreement with extensive experimental data including the detection of long-range CSPs (***Iešmantavičius et al., 2013***), and their recurrence in independent simulations initiated from the bound structure collectively support the conclusion that the integrative OPES ensemble provides new detailed insights into the apo state of ACTR.

## Discussion

IDPs challenge our understanding of structure–function relationships, as their dynamic ensembles populate transient conformations that are often invisible to experiments and difficult to sample in simulations (***Granata et al., 2015***). All-atom explicit-solvent molecular dynamics can provide detailed insights into the structure and dynamics of IDPs, but these insights are only reliable when derived from well-converged ensembles. Such ensembles are essential not only for benchmarking and refining force fields against experimental observables (***Robustelli et al., 2018***; ***Phan et al., 2025***), but also as training data for generative AI-based ensemble predictors (***Lewis et al., 2025***). Here, we show that OPES multithermal simulations efficiently generate converged atomistic ensembles across peptides and IDPs spanning 15–71 residues in length, offering a robust and reproducible framework for exploring IDP conformational landscapes.

We quantified convergence and sampling efficiency using cumulative averages, autocorrelation analyses, error bar comparisons, and two-dimensional projections of the sampled conformational space. For the helical benchmark peptide (AAQAA)_3_, OPES reproduced radius of gyration distributions and per-residue helical propensities in close agreement with REST2 and unbiased MD, achieving 2–3× acceleration relative to REST2 and 10–16× relative to unbiased simulations. For ACTR_20−60_, independently generated biases yielded reproducible ensembles and improved sampling efficiency, while larger systems such as HTTex1 and full-length ACTR explored low-population, highly helical, or compact states that are difficult to access with standard REST2 and unbiased simulations.

OPES combines these sampling advantages with practical simplicity. It eliminates the need for careful optimisation of temperature ladders, allows even a single replica to generate a multithermal ensemble, and enables straightforward reweighting and temperature-dependent analyses. Moreover, OPES naturally accommodates additional collective variable-based biases to further accelerate slow degrees of freedom when needed (***Invernizzi et al., 2020***; ***Bonati et al., 2021***; ***Rizzi et al., 2023***).

Critically, OPES-derived ensembles provide mechanistic insight beyond what experiments alone can reveal. For ACTR, we observed rare yet thermodynamically accessible states in which multiple helices fold cooperatively, forming partially binding-competent conformations (Figure 5). These transiently structured states may reflect an energetic compromise between conformational selection and induced fit (***Kjaergaard et al., 2010a***; ***Iešmantavičius et al., 2014***), in which prestructuring reduces the entropic cost of folding-upon-binding (***Spolar and Record Jr, 1994***; ***Theisen et al., 2021***). Transient tertiary structure has also been suggested for other IDPs such as *α*-synuclein and p53 (***Graen et al., 2018***; ***Szöllősi et al., 2025***), and experiments have detected signatures of bound-like conformations in the apo state ensembles of Gab1 and a dynein IDR (***Morgan et al., 2021***; ***Gruber et al., 2022***). Similar helix preformation has previously also been reported by simulation and experimental studies of the “molten globule”-like NCBD (***Kjaergaard et al., 2010b***; ***Knott and Best, 2012***). Such atomistic resolution insights would be difficult to obtain from experiments alone and simpler coarse-grained models, which lack the resolution needed to capture detailed interactions and cooperative folding events. We anticipate that these atomistic ensembles will be particularly valuable for studies and design of IDP interactions with small molecules (***Heller et al., 2020***; ***Lohr et al., 2022***; ***Zhu et al., 2022***; ***Zhu and Robustelli, 2025***) or other proteins (***Kjaer et al., 2025***), as transiently folded or compact states may harbour specific functional binding pockets and interaction sites.

Despite these advantages, OPES has limitations. System size and simulation box volume impose practical constraints, as increased computational demand is required to converge bias potentials. Narrowing the temperature range may help mitigate this, at the expense of reduced enhanced sampling. Moreover, despite OPES having substantially enhanced conformational exploration relative to REST2 and unbiased MD, it is unlikely to have exhaustively sampled all relevant low-population states, given the formidable sampling challenge posed by these conformationally heterogeneous systems. For example, several conformations observed in MD simulations initiated from the bound state of ACTR were not recovered, or were sampled with reduced structural content. Furthermore, extending OPES to even larger IDPs or multi-protein assemblies will likely require significant computational resources.

Within these constraints, OPES multiT nonetheless provides a robust, simple and reproducible framework for generating atomistic ensembles of IDPs. By complementing experimental data and enabling reproducible sampling of rare, partially folded states at atomic resolution, this approach provides a practical framework for probing IDP dynamics, informing functional studies and drug discovery, guiding disordered protein engineering, and supporting next-generation AI models of structural ensembles.

## Methods

### Molecular dynamics simulations

All simulations were prepared and performed with GROMACS 2023.4 (***Abraham et al., 2015***) and PLUMED 2.10 (***Tribello et al., 2014***) using GPUs (except REST2 simulations, *vide infra*). Table 1 provides an overview of the simulation systems studied in this work including force-field parameters, thermostat temperatures, salt concentrations, and number of water molecules.

All proteins were simulated in a dodecahedral-shaped box, ensuring the box size provides sufficient volume to avoid artifacts related to periodic image protein contacts. Prior to MD, energy minimisation was performed with the steepest descent algorithm. We then used the leap-frog integrator for all MD simulations with a 2 fs timestep. LINCS (***Hess et al., 1997***) was used to constrain all hydrogen bonds. Short-range nonbonded interactions were calculated up to a cut-off of 0.9 and 1.0 nm for simulations with the ff03w/ff03ws and a99SB-disp force fields, respectively. Long-range electrostatic interactions were computed using particle mesh Ewald (PME) (***Darden et al., 1993***). PME calculations were performed with cubic interpolation and a Fourier spacing of 0.12 and 0.125 nm for a03w/a03ws and a99SB-disp force fields, respectively. The velocity rescale method with a time constant of 0.1 ps was used to ensure temperature control (***Bussi et al., 2007***). During the first equilibration in the NPT ensemble, pressure control was achieved with the Berendsen baro-stat (***Berendsen et al., 1984***) with a reference pressure of 1 bar, compressibility of 4.5×10^−5^ bar^−1^ and a coupling constant of 2 ps. For all subsequent NPT simulations we used the Parrinello-Rahman barostat (***Parrinello and Rahman, 1981***).

All systems were initially equilibrated for 500 ps in the NVT ensemble at the target temperature, followed by another equilibration in the NPT ensemble with the Berendsen barostat for 500 ps. These two initial equilibration simulations were performed with position restraints on all protein heavy atoms with a force constant of 1000 kJ mol^−1^ nm^−2^. For unbiased simulations of (AAQAA)_3_ where frames were saved every 100 ps, subsequent production simulations used the Parrinello-Rahman barostat. For OPES and REST2 simulations, we also ran an additional equilibration with the Parrinello-Rahman barostat for 1 ns. We then selected the frame with the closest volume to the average volume calculated from this 1 ns NPT simulation as a starting configuration for all subsequent production simulations in the NVT ensemble.

For all ACTR_20−60_, ACTR, and HTTex1 16Q simulations we then performed an initial simulation at 450 K in the NVT ensemble for at least 100 ns, saving all atomic coordinates every 1 ns. With this step we aimed to sample different structures to be used for subsequent OPES and REST2 production simulations. Thus, for these systems all production simulations described below were initiated from different initial coordinates and random velocities. For (AAQAA)_3_, the same initial coordinates with different random velocities were used due to the smaller size of this peptide.

Unbiased simulations of ACTR from the bound state (residues 23–69, PDB 1KBH, without the NCBD of CBP) were performed after capping the N- and C-terminus with acetyl and N-methyl amide groups, respectively. The a99SB-disp force field and an equilibration protocol identical to all other ACTR simulations were used. Production simulations were then performed in the NPT ensemble (Parrinello-Rahman barostat) at 300 K and 1 bar, saving frames every 100 ps.

### OPES multithermal simulations

In the on-the-fly probability enhanced sampling (OPES) framework, an adaptive bias potential is built on-the-fly to drive the system toward a prescribed target distribution as previously described (***Invernizzi et al., 2020***; ***Trizio et al., 2024***). Here we focus on the expanded ensemble variant of OPES, more precisely on OPES multithermal (multiT). Briefly, the inverse temperature is defined as *β* = (*k*_B_*T*)^−1^, and a reference inverse temperature *β*_0_ is used to propagate the dynamics. In OPES multithermal, the target distribution is a mixture of canonical ensembles over a set of inverse temperatures {*β*_*k*_}:

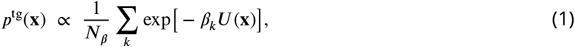

where *U* (**x**) is the potential energy. The adaptive bias at iteration *n* is explicitly

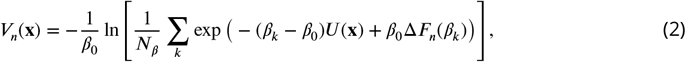

where Δ*F*_*n*_(*β*_*k*_) are on-the-fly estimates of the relative free energies associated with each temperature state. At convergence, the bias becomes quasi-static, enforcing approximately uniform sampling across the temperature range. Once the bias has converged, ensemble averages at any target inverse temperature *β*^∗^ are computed by reweighting the biased frames. For a frame *i* with potential energy *U*_*i*_ and bias *V* (**x**_*i*_), the unnormalised weight is

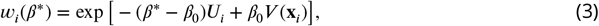

and normalised weights 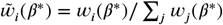 are used to calculate ensemble-averaged values of any property of interest 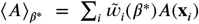. Furthermore, the statistical reliability of the weighted ensemble is quantified by the Kish effective sample size across the temperature range,

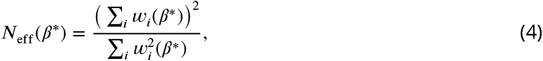

which measures the equivalent number of independent, equally weighted samples.

In practice, for each system we first chose an appropriate number of temperature steps for the OPES target distribution (i.e., temperature range to be sampled, typically 290–450 K). One can estimate the number of temperature steps by running multiple short (500 ps) simulations without specifying the `TEMP_STEPS` argument in the PLUMED input file. PLUMED then attempts to estimate the number of steps required based on the potential energy fluctuations of the system (***Invernizzi et al., 2020***). However, this approach provides variable results for the systems studied here (Table S2). We therefore advise practitioners to obtain multiple estimates from independent starting configurations or velocities. Specifically, we used the minimum value of the obtained range as a guide and upper bound for our simulations. Furthermore, based on our simulation systems studied here, we also estimated a simple relationship between 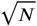 (number of atoms) and the number of temperature steps. For a typical 290–450 K temperature range one can estimate the required number of temperature steps with 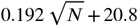 (based on linear regression, Table S2). After determining the number of temperature steps, we initiated ten independent trial simulations in the NVT ensemble to build the bias potential. Trial simulations with limited exploration of temperature (potential energy) space were stopped after 50–100 ns. Simulations exhibiting full coverage of the temperature range were continued until the bias potential converges (Figure S1). To this end, bias convergence can be monitored using

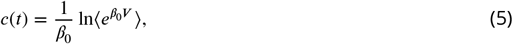

reported by PLUMED. As the bias approaches its quasi-static limit, *c*(*t*) tends toward a constant value, providing an empirical indicator of convergence, although it should not be used for reweighting. Additionally, we calculated sampling scores to quantify coverage of temperature space as the fraction of temperature steps with an *N*_eff_ greater than expected from a uniform distribution (1/N, where N is the number of temperature steps). Values near 1.0 indicate that all temperatures were effectively sampled. The success rates in obtaining converged bias potentials (defined by *c*(*t*) reaching a constant value and a sampling score above 0.8) ranged from 50% ((AAQAA)_3_) to 10% (ACTR) and are shown in Table S2. Alternatively, trial simulations may also be performed with the multiple walker scheme (***Raiteri et al., 2006***), where all trial replicas contribute to the same bias, but we have not explored this here.

Converged bias potentials were then used as static bias potentials (i.e., no further iterative updates to the bias) for production simulations. For larger systems like HTTex1 16Q and ACTR where only a limited number of trial simulations may reach a converged bias, a single static bias may be used for independent production replicas. All production simulation replicas were concatenated as if they originated from a single, continuous run. When analysing concatenated (AAQAA)_3_ simulations with different biases, no extra optimisation was performed to combine the weights beside normalisation, since all bias potentials were on average within 1 *k*_B_*T* of each other. Simulation data (bias potential, potential energy, coordinates) were saved every 10 ps. With our setup, OPES simulations exhibit 30–40% reduced simulation performance compared to unbiased MD due to the enhanced sampling overhead (Table S1), which is moderate considering the acceleration of sampling. In summary, the procedure to obtain OPES multiT ensembles for IDPs is as follows:

#### 1. Estimate temperature steps

Run multiple (e.g., 10) simulations without specifying `TEMP_STEPS` in PLUMED. Use the minimum observed value as a guide/upper bound. Alternatively, use the empirical formula

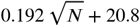

for the 290–450 K temperature range.

#### 2. Bias convergence simulations

Initiate ten independent trial simulations to obtain bias potentials for 50–100 ns. Discard replicas showing minimal exploration of potential energy space, and continue others until the bias converges (monitored by *c*(*t*) and the sampling score).

#### 3. Production simulations

Use one converged bias potential as a static bias. Optionally, start multiple productions with independent biases; these should be very similar when converged.

#### 4. Analysis

Concatenate all production simulations for analysis and account for the OPES bias by reweighting using Eq. 3.

We note that the choice of the integration timestep can affect the accuracy of the calculated potential energy (***Hopkins et al., 2015***), and thus also the bias potential. We therefore recommend using a standard integration timestep of 2 fs with hydrogen bond constraints. Similarly, a 2 fs timestep is expected to lead to decreased accuracy at higher thermostat temperatures, and we therefore recommend using a reference thermostat temperature near the lower end of the temperature range.

To ensure simulation boxes were sufficiently large for all systems studied here we computed the minimum distances between periodic images of protein molecules with the GROMACS mindist tool. None of the simulations exhibited prolonged contacts between periodic images.

For all OPES simulations, we checked whether any peptide bond cis-trans isomerisation events occurred because this may cause some conformational states to become kinetically trapped. With the exception of Pro36 in HTTex1 (transitioned to cis after approx. 1 *μ*s in one replica) and Gly63 in ACTR (which reversibly sampled the cis state for ~150 ns in 1 replica), all peptide bonds remained in the trans configuration at all times. The issues of cis-trans isomerisation and trapped peptide bond conformations are thus not a significant concern with the temperature range and force fields used in this work.

### REST2 simulations

REST2 (***Wang et al., 2011***; ***Bussi, 2014***; ***Koneru et al., 2025***) simulations were performed on CPUs with GROMACS 2022.4 and PLUMED 2.8.2. All systems were simulated using an effective solute temperature range of 300 to 500 K, except HTTex1 16Q, which was simulated in the range of 293 to 500 K. The number of temperature replicas (spaced geometrically in temperature space) and range of exchange probabilities (minimum–maximum) averaged over the entire production simulations are summarised in Table 2. All replicas were initiated from different coordinates and random velocities. The replicas were first equilibrated without replica exchange for 100 ps using the new Hamiltonians with scaled solute temperatures. During production simulations in the NVT ensemble, replica exchange was then attempted every 1.6 ps, saving coordinates every 40 ps. To ensure different initial replicas have sampled similar regions of conformational space, secondary structure and protein R_*g*_ distributions were analysed for temperature replicas (time discontinuous) and demuxed replicas. Demuxed replicas are the time-continuous trajectories reconstructed after the demultiplexing step, in which REST2 trajectories—originally fragmented across replicas due to exchange—are reordered to follow a single physical system continuously in time. Additionally, we also quantified the number of replica round trips in temperature space, average effective solute temperature of each demuxed trajectory, and the fraction of time each demuxed trajectory spent in the ground replica. All ensemble-averaged statistics were derived from the ground replica (original Hamiltonian).

**Table 2.** REST2 replica temperature ranges and exchange probabilities for all systems.

| System | Replicas | Temp. range (K) | Exchange probability |
| --- | --- | --- | --- |
| (AAQAA)<sub>3</sub> | 8 | 300–500 | 38–48 % |
| ACTR<sub>20–60</sub> | 16 | 300–500 | 36–43 % |
| ACTR | 20 | 300–500 | 33–43 % |
| HTTex1 16Q | 20 | 293–500 | 40–49 % |

### Simulation analyses

We used the Python package pyblock^1^ to calculate standard errors of ensemble-averaged quantities. Pyblock implements the blocking analysis described by ***Flyvbjerg and Petersen (1989)***. The same procedure was applied to data from unbiased, REST2, and OPES simulations to obtain error estimates in a consistent manner. Errors obtained from blocking of the concatenated OPES simulations were similar to the standard errors of the mean calculated from averages of individual simulations (Figure S1c). We note, however, that blocking can sometimes underestimate the true uncertainty, as mean and error estimates from individual simulations were not always mutually consistent (Figure S1c). This is not expected to affect the comparison of errors across different simulation methods, since all errors were estimated using the same blocking procedure.

Radii of gyration were calculated using PLUMED (non-mass weighted, utilised for all timecourse and free-energy landscape analyses) and the python library MDTraj for mass-weighted averages comparable to SAXS-derived values (***McGibbon et al., 2015***). To quantify the total helicity of the proteins studied here, we used a helicity parameter based on the `ALPHARMSD` collective variable implemented in PLUMED. The collective variable was computed using optimal rotational and translational alignment (`TYPE=OPTIMAL`) of all possible, continuous six residue fragments against an ideal helical conformation. For each fragment, the RMSD, denoted *r*, was transformed into a smooth indicator function using a rational switching function with parameters *R*_0_ = 0.08 nm, *n* = 8, and *m* = 12. The switching function is defined as

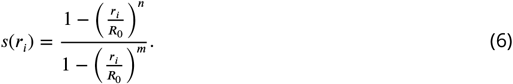

The `helicity` collective variable was then obtained by summing these contributions over all *N* residues,

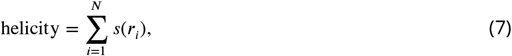

where *N* corresponds to the number of possible fragments. Residue-resolved secondary structure analyses employed DSSP (***Kabsch and Sander, 1983***) implemented in GROMACS.

To analyse transient folding of ACTR compared to reference structures, we calculated the fraction of “native” contacts (Q) with respect to given reference structures. SMOG2 was used to extract native contact pairs with default parameters (***Noel et al., 2016***). MDAnalysis (***Michaud-Agrawal et al., 2011***) and Numpy (***Harris et al., 2020***) were used to extract coordinates and distances for all native contact pairs. Q was then calculated using the switching function proposed by ***Best et al.(2013)***:

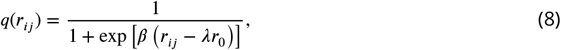

where *r*_0_ denotes the distance in the reference structure, *λ* (set to 1.8) shifts the midpoint of the transition, and *β* (set 50 nm^−1^) controls the sharpness of the switching behavior. These values were then averaged over all contact pairs for each simulation frame. As reference structures, we used chain A of PDB 1KBH (***Demarest et al., 2002***), representing the native conformation bound to CBP. Additionally, we also used cluster centroids obtained from unbiased MD simulations initiated from chain A of PDB 1KBH, representing helical states with tertiary structure that are stable on the microsecond timescale.

Time autocorrelation functions were computed for scalar and vector-valued observables using a fast Fourier transform (FFT)–based approach implemented in Numpy. For a scalar time series *x*(*t*_*k*_) of length *N*, the mean was first subtracted,

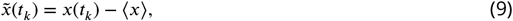

and the normalised autocorrelation function was evaluated via the Wiener-Khinchin theorem as

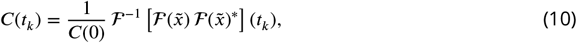

where ℱ and ℱ^−1^ denote the forward and inverse FFT, respectively, and ^∗^ indicates complex conjugation. Zero-padding to length 2*N* was used to avoid circular convolution artifacts. For DSSP time-series (a vector-valued observable) **X**(*t*), the DSSP strings were first converted to number (1=helix, 2=strand, 3=coil), and the autocorrelation function was computed by summing the power spectra of the individual components,

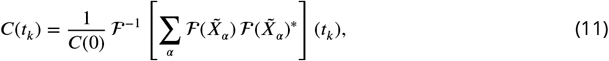

where *α* runs over the components and 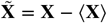. The integrated autocorrelation time *τ*_int_ was estimated as the area under the autocorrelation function,

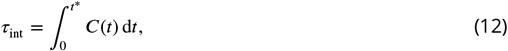

where the upper limit *t*^∗^ was chosen as the first time at which *C*(*t*) becomes negative, thereby reducing the contribution of statistical noise at long times. The integral was evaluated numerically using the trapezoidal rule.

MD convergence was assessed using quantitative stabilisation criteria applied to cumulative ensemble averages and associated uncertainties. For the R_*g*_ and total fraction helix (derived from DSSP), the stabilisation time of the mean was defined as the earliest time at which the relative fluctuation of the cumulative average within a sliding window of five consecutive time points fell below 10%. The relative fluctuation was computed as the peak-to-peak variation within the window normalised by the window-averaged value. Error stabilisation was defined independently as the first time at which the relative error (error divided by the absolute value of the observable) remained below 10% for all points within the same five-point window. For per-residue helical fraction profiles, convergence was evaluated using RMSD-based and error-based thresholds. The helix profile was considered stabilised when the RMSD between the cumulative helix profile and the full-trajectory ensemble profile fell below 2%. Independently, stabilisation of the profile uncertainty was defined as the first time at which the average per-residue error dropped below 2%. In cases where no threshold was reached, the final simulation time was conservatively assigned as the stabilisation time. For both analyses, if either criterion was not met, the final simulation time was reported.

Conformational exploration was quantified by projecting trajectories onto two-dimensional collective variable (CV) spaces and analysing the resulting probability distributions using bootstrap-resampled, effective-sample-size *N*_eff_ –matched ensembles. For OPES simulations, the *N*_eff_ was computed from the OPES reweighting factors (Eq. 4), while REST2 ensembles were subsampled to the same *N*_eff_ to enable unbiased comparisons. Two-dimensional histograms were constructed using a uniform grid, with the number of bins per dimension chosen as 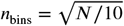, corresponding to an average of approximately ten samples per bin. Three complementary metrics were evaluated on each bootstrap replica (500 iterations): (i) the effective coverage, defined as the normalised effective number of occupied bins based on histogram counts; (ii) the normalised Shannon entropy of the 2D distribution, scaled by the maximum possible entropy for the grid; and (iii) the relative coverage, defined as the fraction of bins occupied in one ensemble that were also sampled by the other. Reported values correspond to bootstrap means and standard deviations. Robustness with respect to histogram resolution was evaluated by repeating the analysis using a range of bin numbers from 0.25 to 1.75 times the default value. We used ELViM with default parameters (***Viegas et al., 2024***) for projecting the ensembles onto a 2D grid. ELViM was run with the merged OPES+REST2 ensembles to ensure projection onto a shared grid. Additionally, we also used 2D projections with the R_*g*_ and helicity parameter as CVs, to better resolve states with different amounts of secondary structure.

Conformational clustering of ACTR simulations and subensembles was performed using the GROMACS cluster tool. We used backbone atoms of residues 27–63 for alignment to calculate the RMSD as a similarity measure combined with a 0.4 nm cut-off for clustering with the gromos method (***Daura et al., 1999***).

### Bayesian/maximum entropy reweighting

We used the ensembles of ACTR produced by our REST2 and OPES multiT simulations as prior ensembles for ensemble reweighting with experimental data using a Bayesian/maximum entropy approach (***Bottaro et al., 2020***). The REST2 ground replica (300 K) and an OPES frames with non-zero weights at 310 K (w > 10^−10^) were used. In both cases, we initialised the reweighting calculations with uniform prior weights. OPES-derived weights yield ensembles and experimental observables that are highly similar to those obtained with uniformly averaged OPES frames, but they introduce a non-uniform baseline for reweighting, making the changes in *N*_eff_ upon subsequent reweighting not directly comparable to the REST2 reweighting calculations. NMR chemical shifts, residual dipolar couplings (RDCs) and paramagnetic relaxation enhancement (PRE) measurements have all previously been measured at 310 K (***Iešmantavičius et al., 2013***). We also used a previously published SAXS dataset measured at 318 K (***Kjaergaard et al., 2010a***). Chemical shifts were calculated using SPARTA+ (***Shen and Bax, 2010***) and secondary chemical shifts were extracted using random coil values predicted with POTENCI (***Nielsen and Mulder, 2018***). RDCs were calculated with the PALES software using a local alignment window of 15 residues (***Zweckstetter, 2008***). PREs were calculated using a rotamer libraries of the MTSL spin label implemented in the DEER-PREdict software (***Tesei et al., 2021***). We assumed a transverse relaxation rate of 10 s^−1^ in the diamagnetic state and an effective spin label correlation time, *τ*_*t*_, of 0.1 ns. SAXS scattering curves were calculated using Pepsi-SAXS (***Grudinin et al., 2017***) with fixed parameters for the hydration layer (*δρ*) and the displaced solvent (*r0*) previously determined (***Pesce and Lindorff-Larsen, 2021***).

For chemical shifts we compared calculated and experimental values using a normalised RMSE (equivalent to the square root of the reduced chi-squared 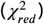:

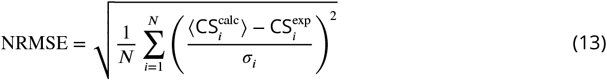

where *σ*_*i*_ is the uncertainty of the calculated chemical shift due to the forward model (nucleus-specific). Additionally, we scaled these errors by the residue-averaged disorder g-score, which quantifies the extent of disorder (g-score = 0) versus order (g-score = 1) (***Invernizzi et al., 2025***). For RDCs, we scaled the magnitude of the ensemble-averaged values to optimise the agreement with experimental data, using residues whose predicted RDCs matched the sign of the experimental values (***Schnapka et al., 2025***). We used the Q-factor calculated with the scaled RDCs to quantify agreement with the experimental data:

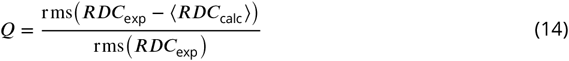

For PREs, we quantified agreement with experimental data by calculating the RMSE of the calculated and experimental intensity ratios 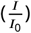. The correlation time of the nucleus-electron vector, *τ*_*C*_, was scanned every 1 ns between 1 and 20 ns to find the value yielding optimal agreement with the experimental data. The calculated SAXS scattering curves were globally scaled to the experimental values with a constant background by least-square fitting using Scikit-learn (***Pedregosa et al., 2011***). Agreement was then quantified using the reduced *χ*^2^:

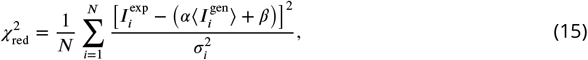

The errors of the SAXS experiments were rescaled using the Bayesian Indirect Fourier Transform (BIFT) method (***Larsen and Pedersen, 2021***).

All ensemble reweighting calculations were performed with the BME software (***Bottaro et al., 2020***). We performed two types of reweighting calculations, one integrating the CS+SAXS data and another with CS+PRE restraints. For both, we scaled the errors of the two types of data (CS+SAXS or CS+PRE) to balance the relative contributions of CS and SAXS/PRE restraints. The experimental uncertainties were rescaled such that the total inverse-variance weight of both datasets was equal. These rescaled errors were only utilised internally for the optimisation calculations and the final reduced *χ*^2^ was calculated with the original errors. An iterative reweighting procedure (***Pesce and Lindorff-Larsen, 2021***) was used to optimise ensemble weights and SAXS scaling and offset constants (*α* and *β*).

For CS+PRE reweighting, PRE data were integrated by implementing pseudo-restraints. Experimental PRE intensity ratios were numerically converted to transverse PRE rates (Γ_2_), which can be linearly averaged, while simulated Γ_2_ values were computed per frame using precalculated distance and angular terms with a fixed rotational correlation time, *τ*_*c*_. Frames for which PRE observables could not be evaluated for all sites (due to steric clashes between spin label rotamers and protein atoms) were excluded from the analysis. Experimental PRE restraints were treated as equality or inequality constraints depending on the intensity ratio, with upper and lower bounds applied for *I*/*I*_0_ > 0.8 and *I*/*I*_0_ < 0.2, respectively. To balance the relative information content of CS and PRE data, experimental uncertainties were rescaled such that the total inverse-variance weight of both datasets was equal. This reweighting procedure was repeated for *τ*_*c*_ values in the range 1–10 ns, thereby treating *τ*_*c*_ as a nuisance parameter. All reweighting calculations were cross-validated with left-out datasets (RDCs and either PRE or SAXS), while keeping the effective fraction of frames, *ϕ*,

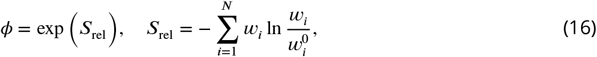

above 0.2.

## Data availability

All OPES multiT MD simulation data and the reweighted ACTR ensemble produced in this work are available on Zenodo at https://zenodo.org/records/18249727. The unbiased simulations, REST2 simulations and ELViM projection data are available on Zenodo at https://zenodo.org/records/18253563. The previously reported unbiased MD trajectory of human ACTR (***Robustelli et al., 2018***) is available for non-commercial use by request from D. E. Shaw Research.

## Code availability

Jupyter notebooks containing Python code used for all analyses presented in this work are available on GitHub at https://github.com/KULL-Centre/_2026_streit_opes. PLUMED input and analysis files for the OPES simulations are available on PLUMED-NEST at https://www.plumed-nest.org/eggs/26/000/.

## Acknowledgments

This work was sponsored by Peptone Ltd. J.O.S. acknowledges support by the European Molecular Biology Organisation through Postdoctoral Fellowship grant ALTF 98-2025. We acknowledge access to computational resources from the ROBUST Resource for Biomolecular Simulations (supported by the Novo Nordisk Foundation grant no. NNF18OC0032608) and via a grant from the Carlsberg Foundation (CF21-0392). We also thank John Christodoulou and the UK High-End Computing Consortium for Biomolecular Simulation, HECBioSim (http://hecbiosim.ac.uk), supported by EPSRC (grant no. EP/X035603/1), for access to ARCHER2 (https://www.archer2.ac.uk). We thank D. E. Shaw Research for sharing the previously reported (***Robustelli et al., 2018***) MD trajectory of human ACTR.

## Author contributions

M.I., S.B. and K.L.-L. conceived the study and all authors contributed to its design. K.T. and K.L.-L. sourced the funding. J.O.S. performed and analysed all simulations with guidance from M.I., S.B. and K.L.-L. J.O.S. wrote the first draft of the manuscript with input from all authors. All authors contributed to the writing and editing of the manuscript.

## Footnotes

1 https://github.com/jsspencer/pyblock

