## Supporting figures and tables for "Transient tertiary structure in intrinsically disordered proteins revealed by multithermal enhanced sampling"

### Supplementary information

#### Supplementary figures

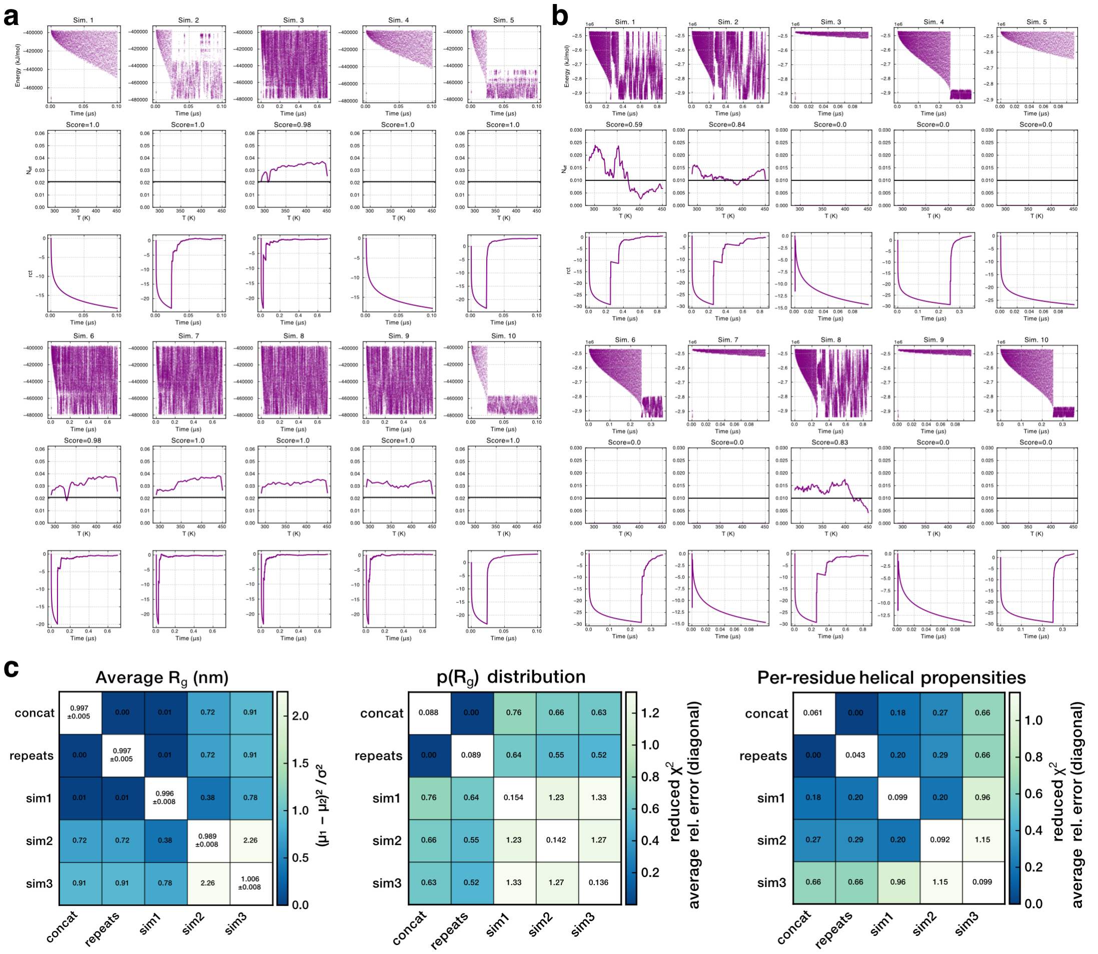


**Figure S1.** Exemplar OPES multiT trial simulations to converge bias potentials for production simulations and comparison of statistical errors obtained from independent production runs and concatenated data. Trial simulations are shown for **a.** $(AAQAA)_3$ at 300 K and **b.** HTTex1 16Q. For each trial simulation, the top row shows the potential energy fluctuations over time, the middle rows the effective sample size across temperatures, and the bottom plot assesses the convergence of the bias potential as a function of time. To calculate effective sample sizes, the first 100 ns, 200 ns and 400 ns were discarded for $(AAQAA)_3$, $ACTR_{20-60}$/ACTR and HTTex1, respectively. The converged bias potentials with the highest scores (fraction of temperature steps with an $N_{\text{eff}}$ above 1/N, where N is the number of temperature steps) were selected for production simulations with a constant bias. **c.** Comparisons of statistical errors obtained for $(AAQAA)_3$ from three production simulations with independent bias potentials (Figure S2b) for the average $R_g$, $R_g$ probability distributions and per-residue helical propensity profiles all at 300 K. Data were either concatenated and errors obtained by blocking (see Methods, concat), the standard error of the mean was obtained from the three replicates (repeats) or errors were calculated for the individual simulations (sim1–3) using blocking. The diagonal values show the average $R_g$ and their associated errors and the average relative uncertainty for the $R_g$ distributions and helical profiles. Agreement between different averages and their combined uncertainties were quantified with the reduced $\chi^2$, with values below 1 indicating estimates are within error of each other.

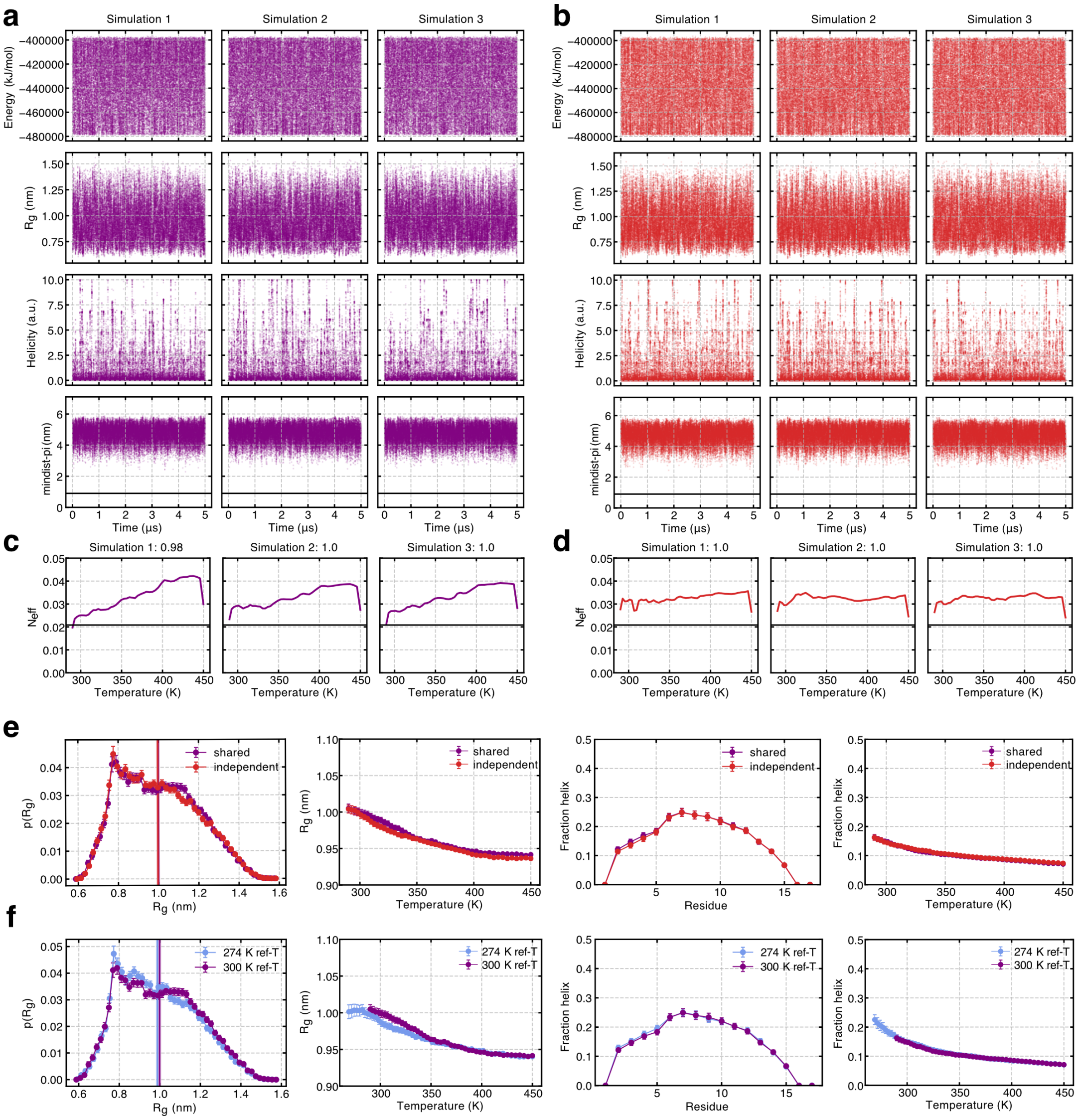


**Figure S2.** OPES multiT simulations of $(AAQAA)_3$ with different bias potentials at a reference temperature of 300 K. **a-b.** Fluctuations of the system potential energy, peptide radius of gyration ($R_g$), total helicity, and minimum distance to a periodic image sampled with OPES multithermal simulations with a shared, constant bias potential (panel a) and independent, constant bias potentials (panel b). **c-d.** Effective sample sizes across the range of temperatures sampled in the simulations from panels a-b. Sampling scores in the subplot titles represent the fraction of temperature steps with an $N_{\mathrm{eff}}$ greater than expected from a uniform target distribution. **e.** Probability distribution of the $R_g$ at 300 K, average $R_g$ values as function of temperature, per-residue helical fractions at 300 K, and total helical fraction as a function of temperature. Results are shown for both the simulation set using a shared and multiple independent OPES biases. **f.** Comparison of ensemble-averaged properties from panel e with OPES multiT simulations conducted at a reference temperature of 274 K and reweighted at 300 K.

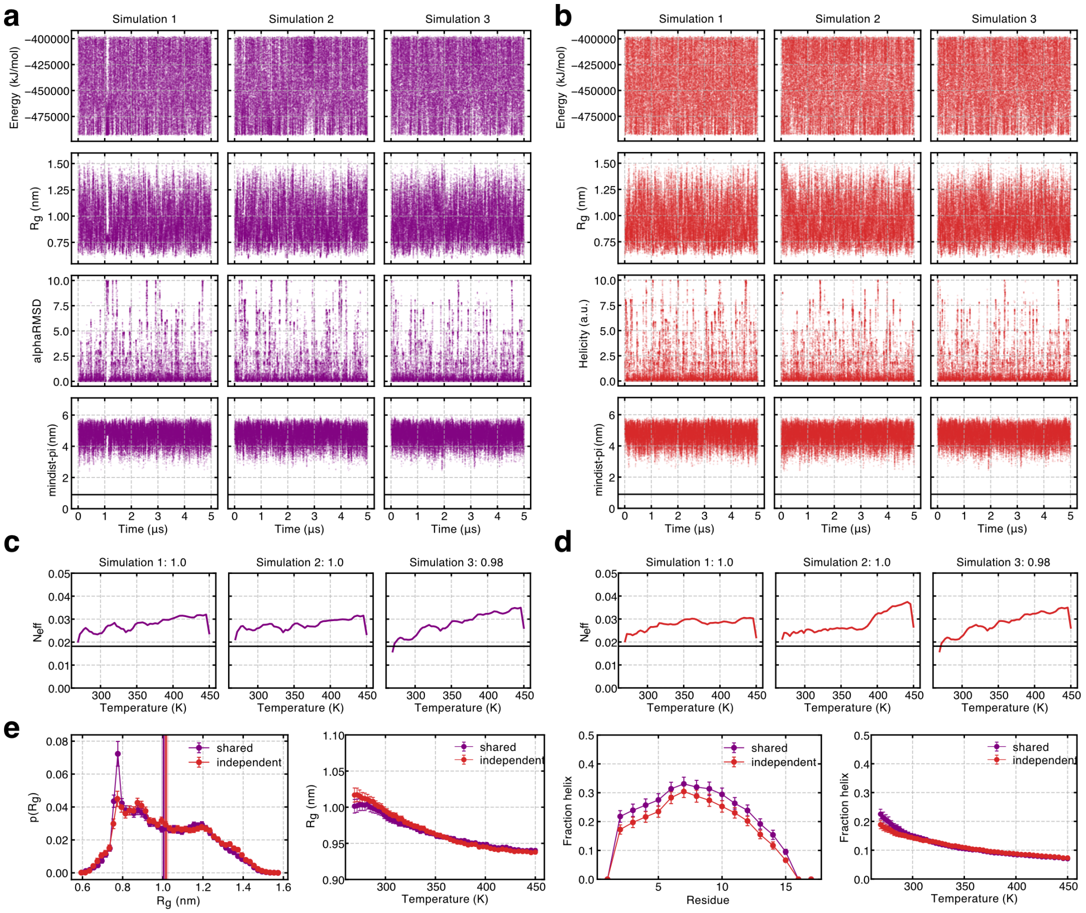


**Figure S3.** OPES multiT simulations of $(AAQAA)_3$ with different bias potentials at a reference temperature of 274 K. **a-b.** Fluctuations of the system potential energy, peptide $R_g$, total helicity, and minimum distance to a periodic image sampled with OPES multithermal simulations with a shared, constant bias potential (panel a) and independent, constant bias potentials (panel b). **c-d.** Effective sample sizes and sampling scores across the range of temperatures sampled in the simulations from panels a-b. **e.** Probability distribution of the $R_g$ at 274 K, average $R_g$ values as function of temperature, per-residue helical fractions at 274 K, and total helical fraction as a function of temperature. Results are shown for both the simulation set using a shared and multiple independent OPES biases.

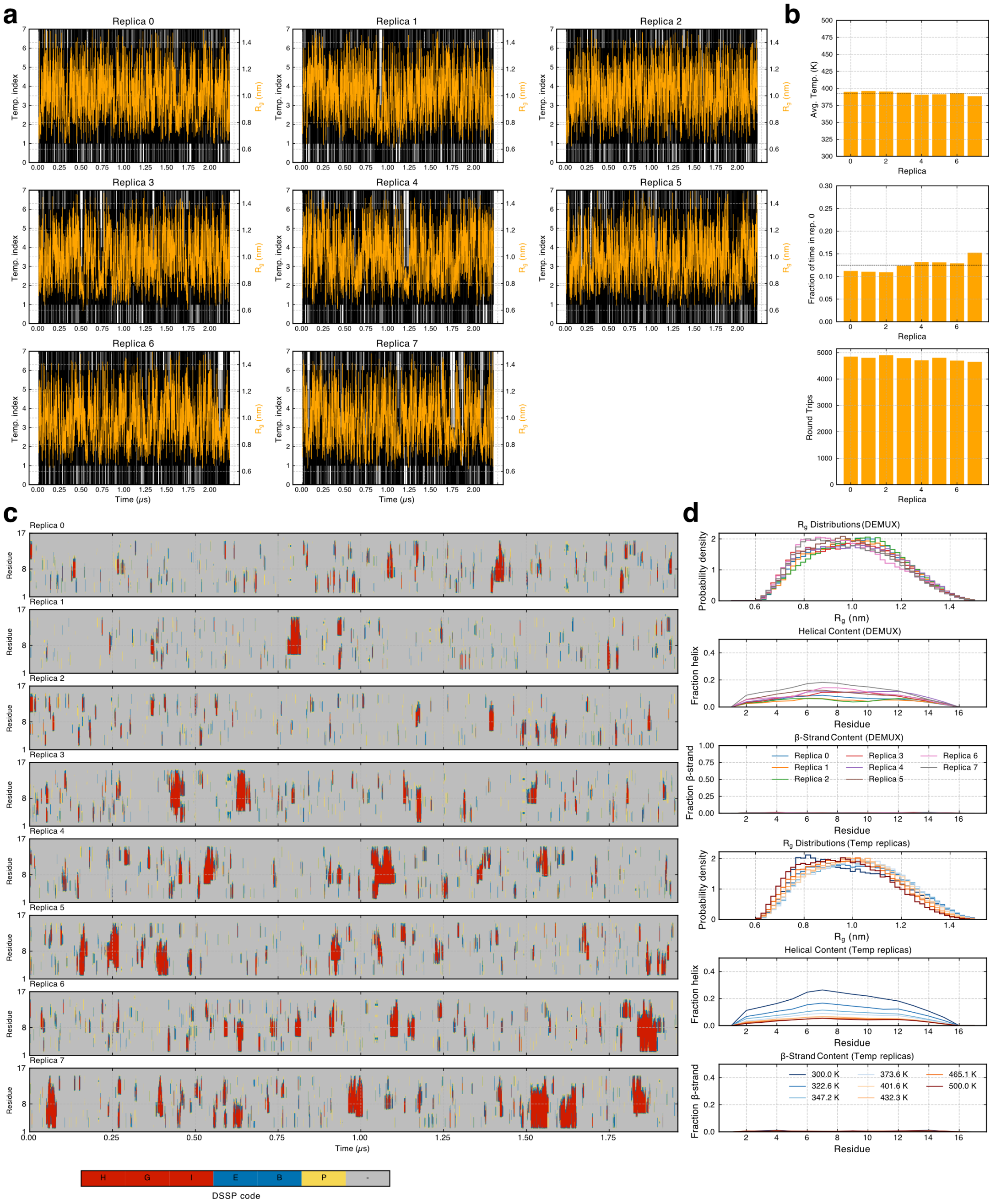

**Figure S4.** REST2 simulations of $(AAQAA)_3$. **a.** Demuxed time series of all replicas showing their traversal through temperature replicas (left axis, black) and fluctuations of the $R_g$ (right axis, orange). **b.** Assessment of REST2 replica mixing. Top: average temperature of each demuxed replica over the course of the entire simulation. Middle: fraction of time spent in the lowest temperature replica. Bottom: number of round trips through temperature space. **c.** Time-resolved secondary structure of the peptide calculated from demuxed trajectories and coloured according to per-residue DSSP assignment. **d.** $R_g$ distributions, per-residue helical fractions and per-residue $\beta$-strand fractions (top to bottom, respectively) calculated from demuxed and temperature replicas.

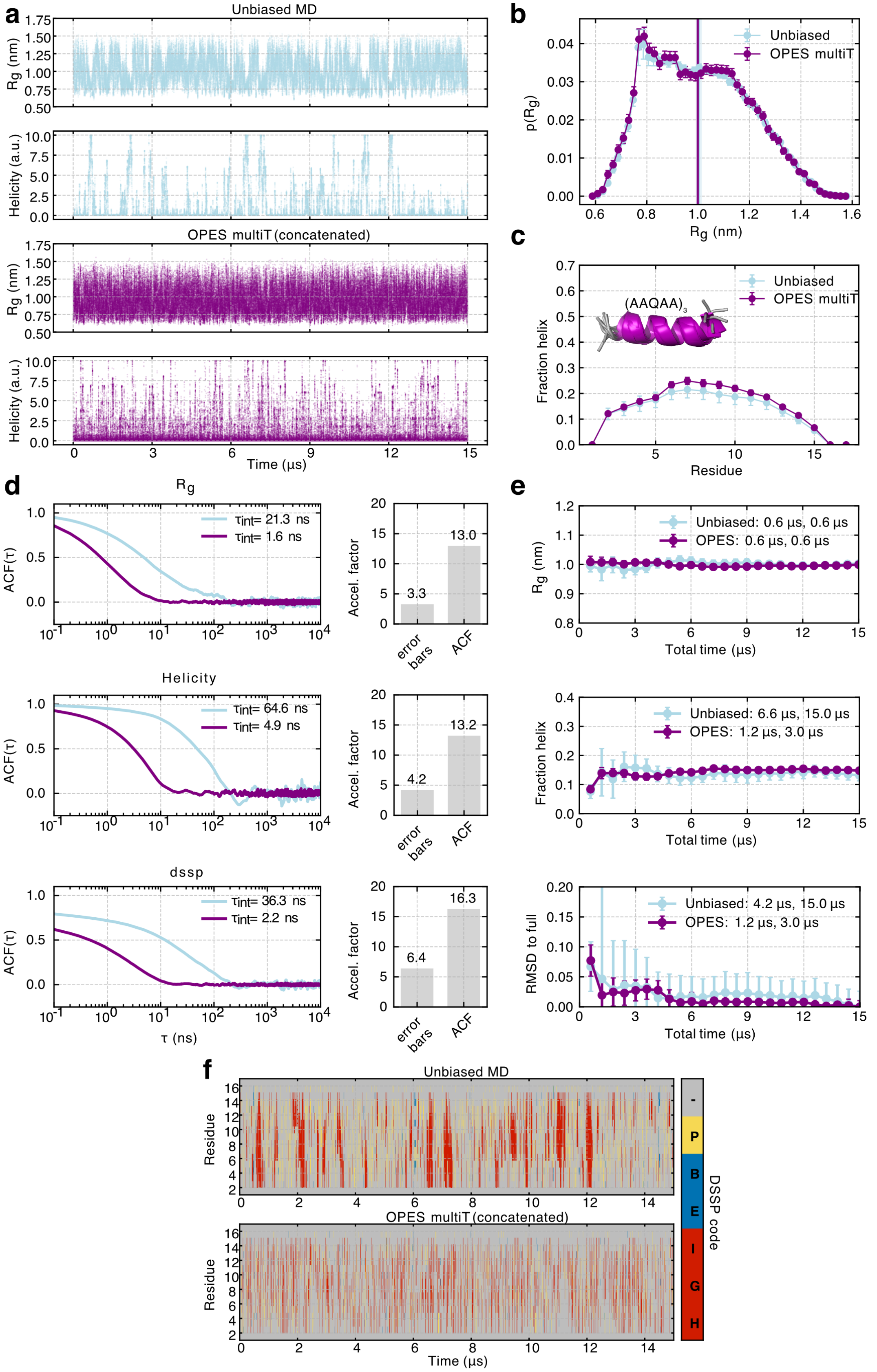
a
Unbiased MD
Rg (nm)
Helicity (a.u.)
OPES multiT (concatenated)
Time (μs)
b
Unbiased
OPES multiT
p(Rg)
Rg (nm)
c
(AAQAA)3
Fraction helix
Residue
d
Rg
τint= 21.3 ns
τint= 1.6 ns
ACF(τ)
Accel. factor
3.3
13.0
error bars
ACF
Helicity
τint= 64.6 ns
τint= 4.9 ns
4.2
13.2
dssp
τint= 36.3 ns
τint= 2.2 ns
6.4
16.3
τ (ns)
e
Unbiased: 0.6 μs, 0.6 μs
OPES: 0.6 μs, 0.6 μs
Total time (μs)
Unbiased: 6.6 μs, 15.0 μs
OPES: 1.2 μs, 3.0 μs
Unbiased: 4.2 μs, 15.0 μs
OPES: 1.2 μs, 3.0 μs
RMSD to full
f
Unbiased MD
OPES multiT (concatenated)
DSSP code
-
P
B
E
I
G
H

**Figure S5.** Comparison of OPES multiT with unbiased (AAQAA)$_3$ simulations at 300 K. **a.** Fluctuations of the radius of gyration (R$_g$) and peptide helicity sampled with unbiased and OPES multithermal simulations. **b.** Probability distributions and ensemble-averaged values (vertical lines) of the R$_g$. Errors represent the standard error of the mean calculated with a blocking analysis **c.** Mean and standard errors of per-residue helical fractions. **d.** Normalised autocorrelation function (ACF) of the R$_g$ (top), helicity (bottom) and per-residue secondary structure (dssp) calculated from the unbiased and OPES multiT timeseries sampled every 100 ps. The correlation times were estimated using the integral of the ACF ($\tau_{int}$). The right panels show estimates of an acceleration factor (OPES / unbiased) estimated from the error bars of ensemble-averaged values and correlation times (ACF). **e.** Time-cumulative averages of the R$_g$, total fraction helix and per-residue helical fractions (measuring the RMSD to the profile derived from the full ensemble) are shown together with the extracted stabilisation times, which mark the earliest points at which the running averages and their uncertainties, respectively, converge to stable values within predefined tolerances. **f.** Time-resolved secondary structure of the peptide coloured according to per-residue DSSP assignment.

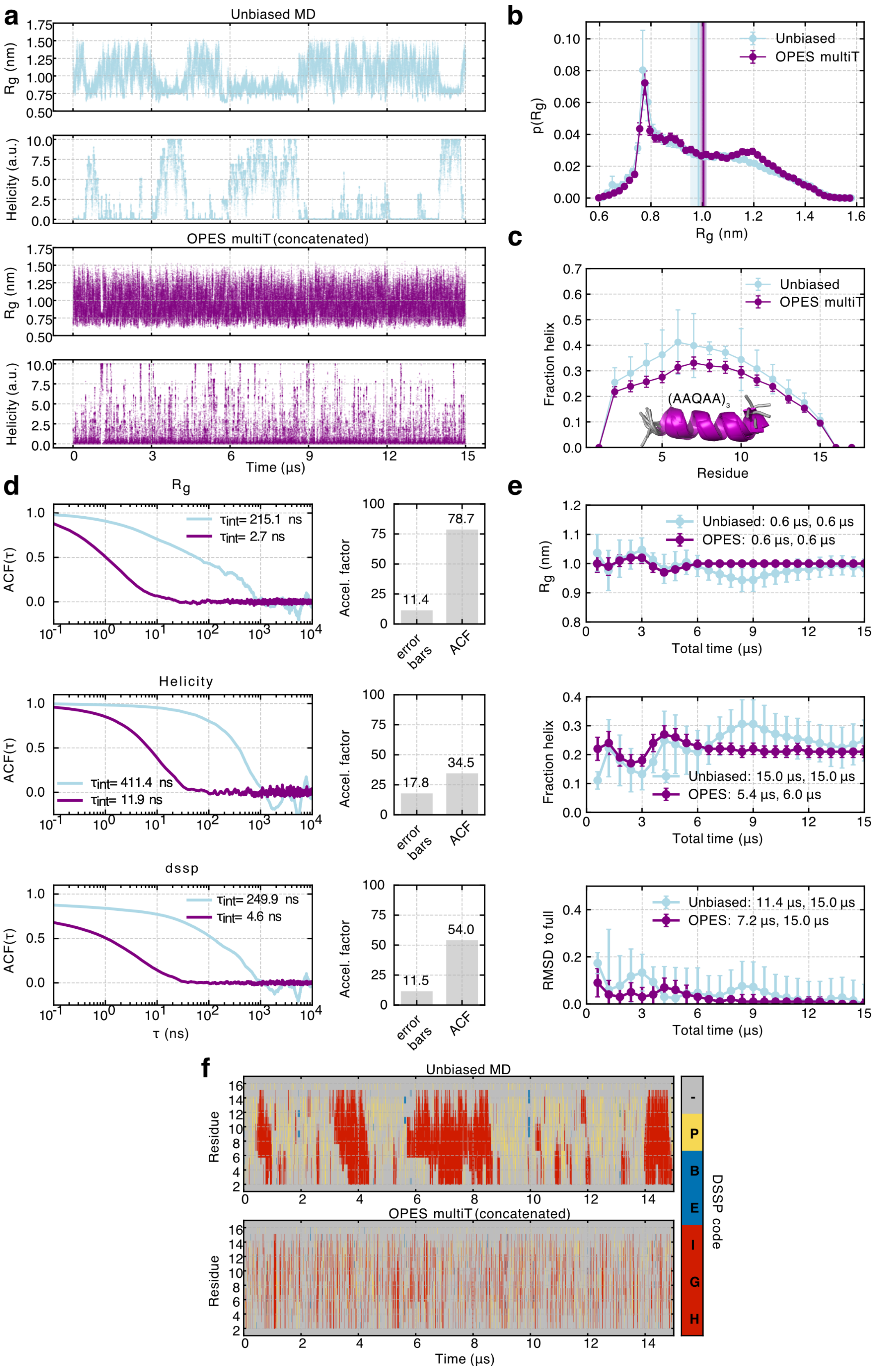

a
Unbiased MD
Rg (nm)
Helicity (a.u.)
OPES multiT (concatenated)
Time (µs)
b
Unbiased
OPES multiT
p(Rg)
Rg (nm)
c
Fraction helix
(AAQAA)3
Residue
d
Rg
ACF(τ)
τint= 215.1 ns
τint= 2.7 ns
Accel. factor
11.4
78.7
error bars
ACF
Helicity
τint= 411.4 ns
τint= 11.9 ns
17.8
34.5
dssp
τint= 249.9 ns
τint= 4.6 ns
11.5
54.0
τ (ns)
e
Unbiased: 0.6 µs, 0.6 µs
OPES: 0.6 µs, 0.6 µs
Total time (µs)
Unbiased: 15.0 µs, 15.0 µs
OPES: 5.4 µs, 6.0 µs
RMSD to full
Unbiased: 11.4 µs, 15.0 µs
OPES: 7.2 µs, 15.0 µs
f
Unbiased MD
OPES multiT (concatenated)
Residue
Time (µs)
DSSP code

**Figure S6.** Comparison of OPES multiT with unbiased $(AAQAA)_3$ simulations at 274 K. **a.** Fluctuations of the radius of gyration ($R_g$) and peptide helicity sampled with unbiased and OPES multithermal simulations. **b.** Probability distributions and ensemble-averaged values (vertical lines) of the $R_g$. Errors represent the standard error of the mean calculated with a blocking analysis **c.** Mean and standard errors of per-residue helical fractions. **d.** Normalised autocorrelation function (ACF) of the $R_g$ (top), helicity (bottom) and per-residue secondary structure (dssp) calculated from the unbiased and OPES multiT timeseries sampled every 100 ps. The correlation times were estimated using the integral of the ACF ($\tau_{int}$). The right panels show estimates of an acceleration factor (OPES / unbiased) estimated from the error bars of ensemble-averaged values and correlation times (ACF). **e.** Time-cumulative averages of the $R_g$, total fraction helix and per-residue helical fractions (measuring the RMSD to the profile derived from the full ensemble) are shown together with the extracted stabilisation times, which mark the earliest points at which the running averages and their uncertainties, respectively, converge to stable values within predefined tolerances. **f.** Time-resolved secondary structure of the peptide coloured according to per-residue DSSP assignment.

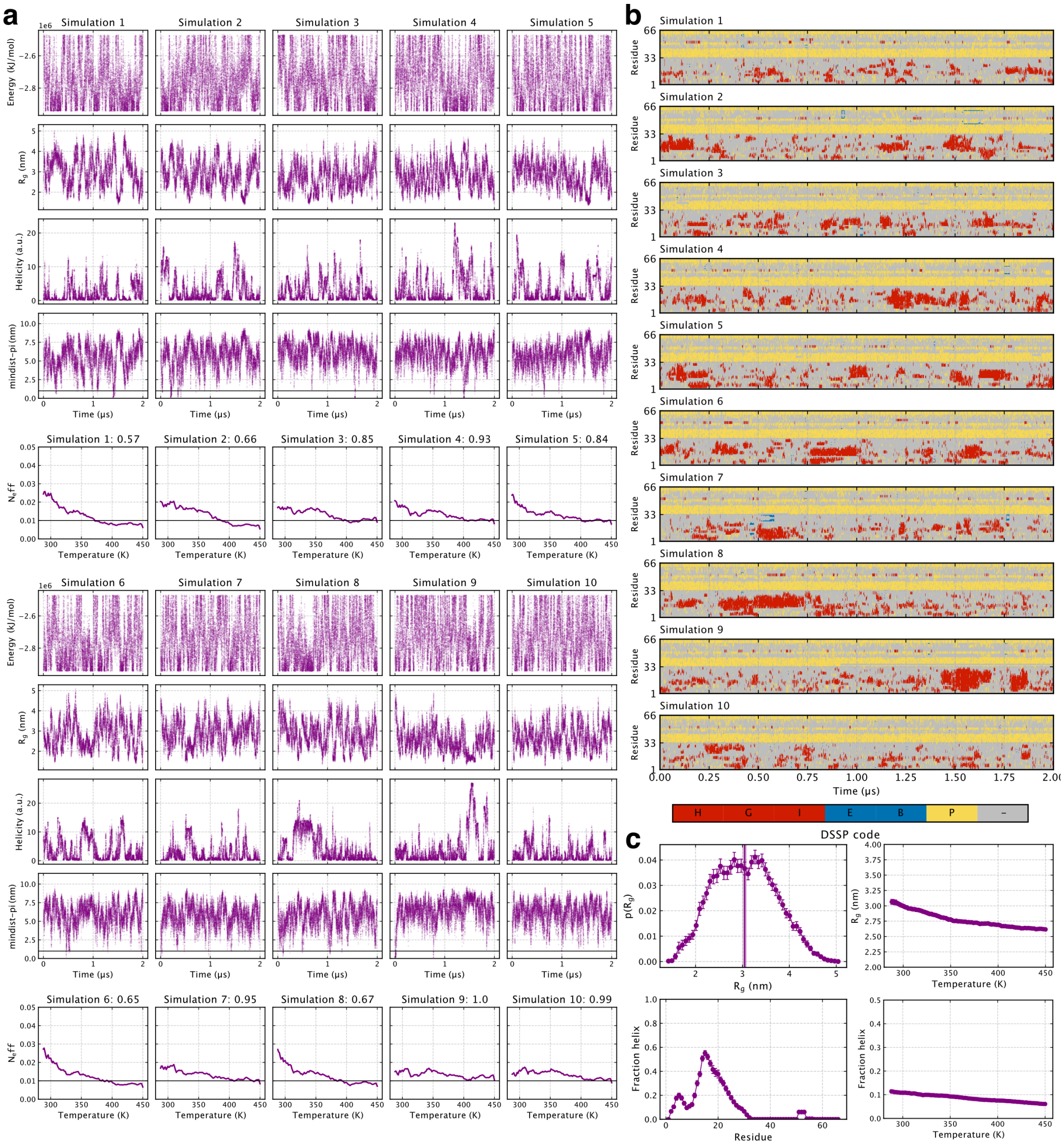
a
Simulation 1
Simulation 2
Simulation 3
Simulation 4
Simulation 5
Energy (kJ/mol)
Rg (nm)
Helicity (a.u.)
mindist-pi (nm)
Time (μs)
Simulation 1: 0.57
Simulation 2: 0.66
Simulation 3: 0.85
Simulation 4: 0.93
Simulation 5: 0.84
Neff
Temperature (K)
Simulation 6
Simulation 7
Simulation 8
Simulation 9
Simulation 10
Simulation 6: 0.65
Simulation 7: 0.95
Simulation 8: 0.67
Simulation 9: 1.0
Simulation 10: 0.99
b
Residue
Time (μs)
H
G
I
E
B
P
–
DSSP code
c
p(Rg)
Rg (nm)
Fraction helix
Residue
Temperature (K)

**Figure S7.** OPES simulations of HTTex1 16Q. **a.** Fluctuations of the system potential energy, protein radius of gyration ($R_g$), total helicity, and minimum distance to a periodic image sampled with OPES multithermal simulations and one shared, static bias potential. Effective sample sizes across the temperature range and sampling scores are shown below each simulation replica. **b.** Time-resolved secondary structure of the protein coloured according to per-residue DSSP assignment. **c.** Probability distribution of the $R_g$ at 293 K, average $R_g$ values as function of temperature, per-residue helical fractions at 293 K, and total helical fraction as a function of temperature.

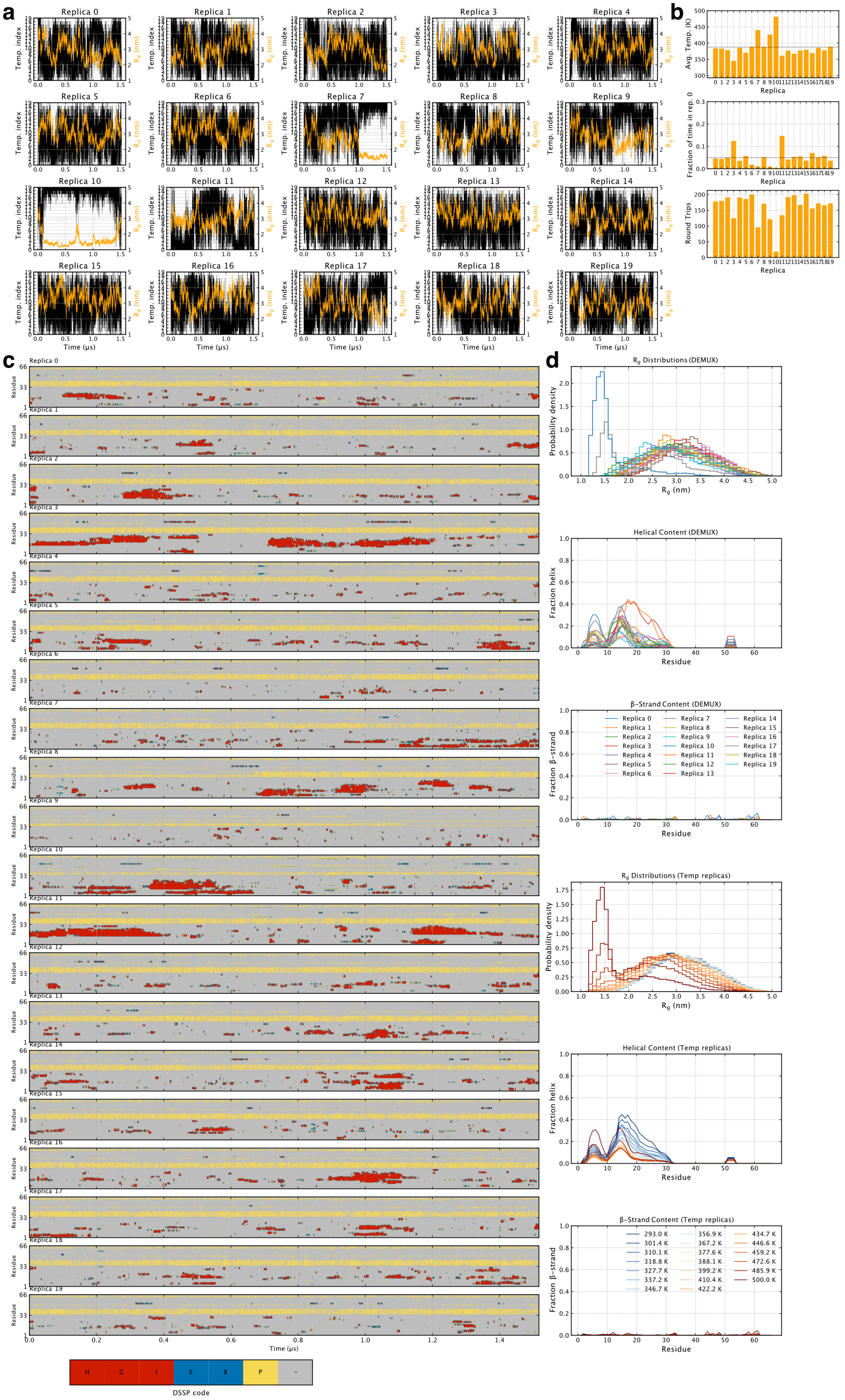
a
b
c
d
Replica 0
Replica 1
Replica 2
Replica 3
Replica 4
Replica 5
Replica 6
Replica 7
Replica 8
Replica 9
Replica 10
Replica 11
Replica 12
Replica 13
Replica 14
Replica 15
Replica 16
Replica 17
Replica 18
Replica 19
Temp. index
Rg (nm)
Time (μs)
Avg. Temp. (K)
Fraction of time in rep. 0
Round Trips
Replica
Residue
DSSP code
H G I E B P ~
Rg Distributions (DEMUX)
Probability density
Rg (nm)
Helical Content (DEMUX)
Fraction helix
β-Strand Content (DEMUX)
Fraction β-strand
Rg Distributions (Temp replicas)
Helical Content (Temp replicas)
β-Strand Content (Temp replicas)
293.0 K
301.4 K
310.1 K
318.8 K
327.7 K
337.2 K
346.7 K
356.9 K
367.2 K
377.6 K
388.1 K
399.2 K
410.4 K
422.2 K
434.7 K
446.6 K
459.2 K
472.6 K
485.9 K
500.0 K

**Figure S8.** REST2 simulations of HTTex1 16Q. **a.** Demuxed time series of all replicas showing their traversal through temperature replicas (left axis, black) and fluctuations of the $R_g$ (right axis, orange). **b.** Assessment of REST2 replica mixing. Top: average temperature of each demuxed replica over the course of the entire simulation. Middle: fraction of time spent in the lowest temperature replica. Bottom: number of round trips through temperature space. **c.** Time-resolved secondary structure of the protein calculated from demuxed trajectories and coloured according to per-residue DSSP assignment. **d.** $R_g$ distributions, per-residue helical fractions and per-residue $\beta$-strand fractions (top to bottom, respectively) calculated from demuxed and temperature replicas.

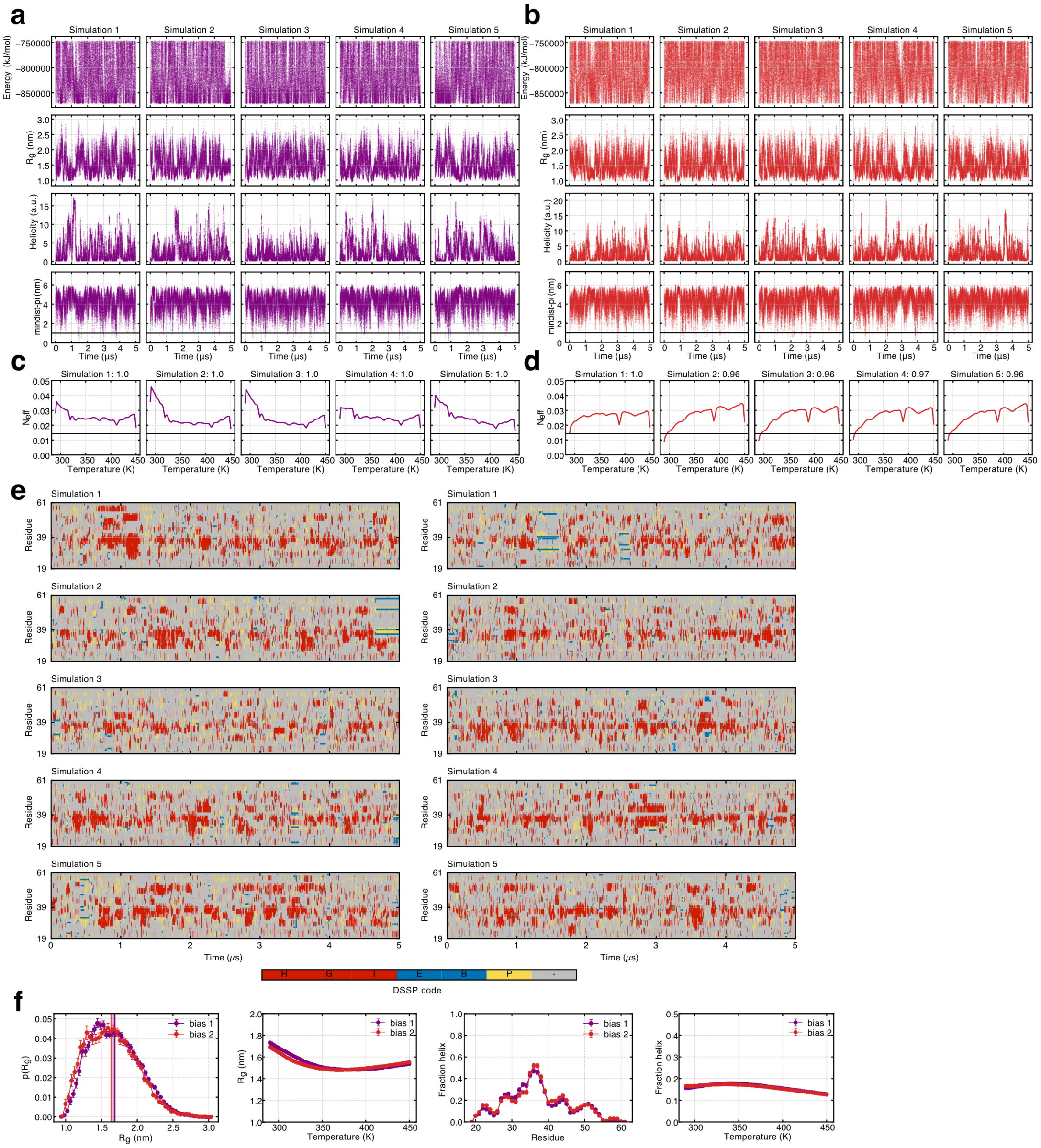

**Figure S9.** OPES simulations of $ACTR_{20-60}$ with different bias potentials. **a-b.** Fluctuations of the system potential energy, protein radius of gyration ($R_g$), total helicity, and minimum distance to a periodic image sampled with OPES multithermal simulations and two independent, static bias potentials. Panel a shows simulations using bias potential 1 and panel b with bias potential 2. **c.** Effective sample sizes across the range of temperatures sampled in the simulations and sampling scores from panels a-b. **e.** Time-resolved secondary structure of the protein simulated with bias 1 (left) and 2 (right) coloured according to per-residue DSSP assignment. **f.** Probability distribution of the $R_g$ at 300 K, average $R_g$ values as function of temperature, per-residue helical fractions at 300 K, and total helical fraction as a function of temperature. Results are shown for both biases.

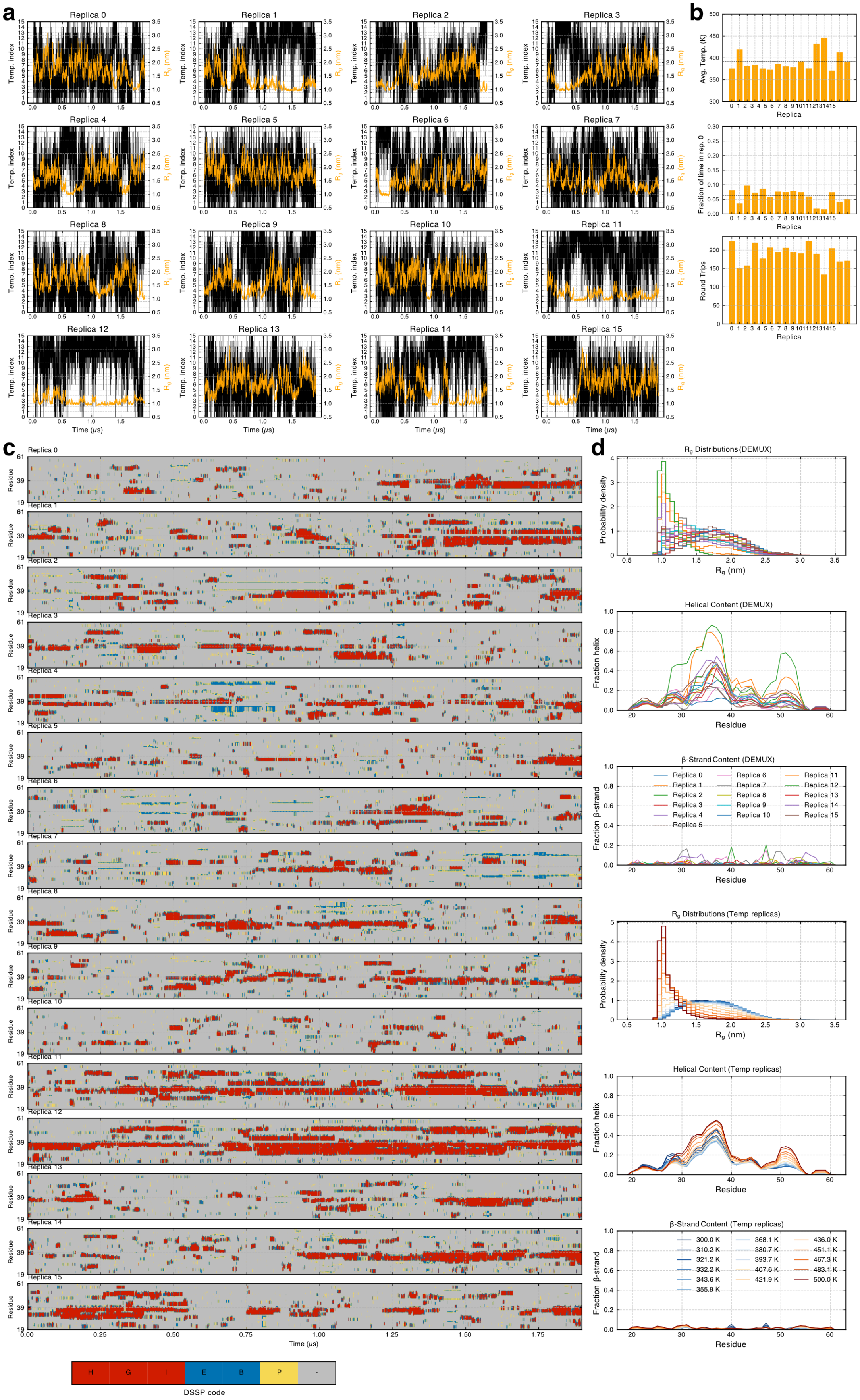
a
Replica 0
Replica 1
Replica 2
Replica 3
Replica 4
Replica 5
Replica 6
Replica 7
Replica 8
Replica 9
Replica 10
Replica 11
Replica 12
Replica 13
Replica 14
Replica 15
Temp. index
Rg (nm)
Time (μs)
b
Avg. Temp. (K)
Fraction of time in rep. 0
Round Trips
Replica
c
Residue
Time (μs)
H
G
I
E
B
P
-
DSSP code
d
Rg Distributions (DEMUX)
Probability density
Rg (nm)
Helical Content (DEMUX)
Fraction helix
Residue
β-Strand Content (DEMUX)
Fraction β-strand
Rg Distributions (Temp replicas)
Helical Content (Temp replicas)
β-Strand Content (Temp replicas)
300.0 K
310.2 K
321.2 K
332.2 K
343.6 K
355.9 K
368.1 K
380.7 K
393.7 K
407.6 K
421.9 K
436.0 K
451.1 K
467.3 K
483.1 K
500.0 K

**Figure S10.** REST2 simulations of ACTR$_{20-60}$. **a.** Demuxed time series of all replicas showing their traversal through temperature replicas (left axis, black) and fluctuations of the R$_g$ (right axis, orange). **b.** Assessment of REST2 replica mixing. Top: average temperature of each demuxed replica over the course of the entire simulation. Middle: fraction of time spent in the lowest temperature replica. Bottom: number of round trips through temperature space. **c.** Time-resolved secondary structure of the protein calculated from demuxed trajectories and coloured according to per-residue DSSP assignment. **d.** R$_g$ distributions, per-residue helical fractions and per-residue $\beta$-strand fractions (top to bottom, respectively) calculated from demuxed and temperature replicas.

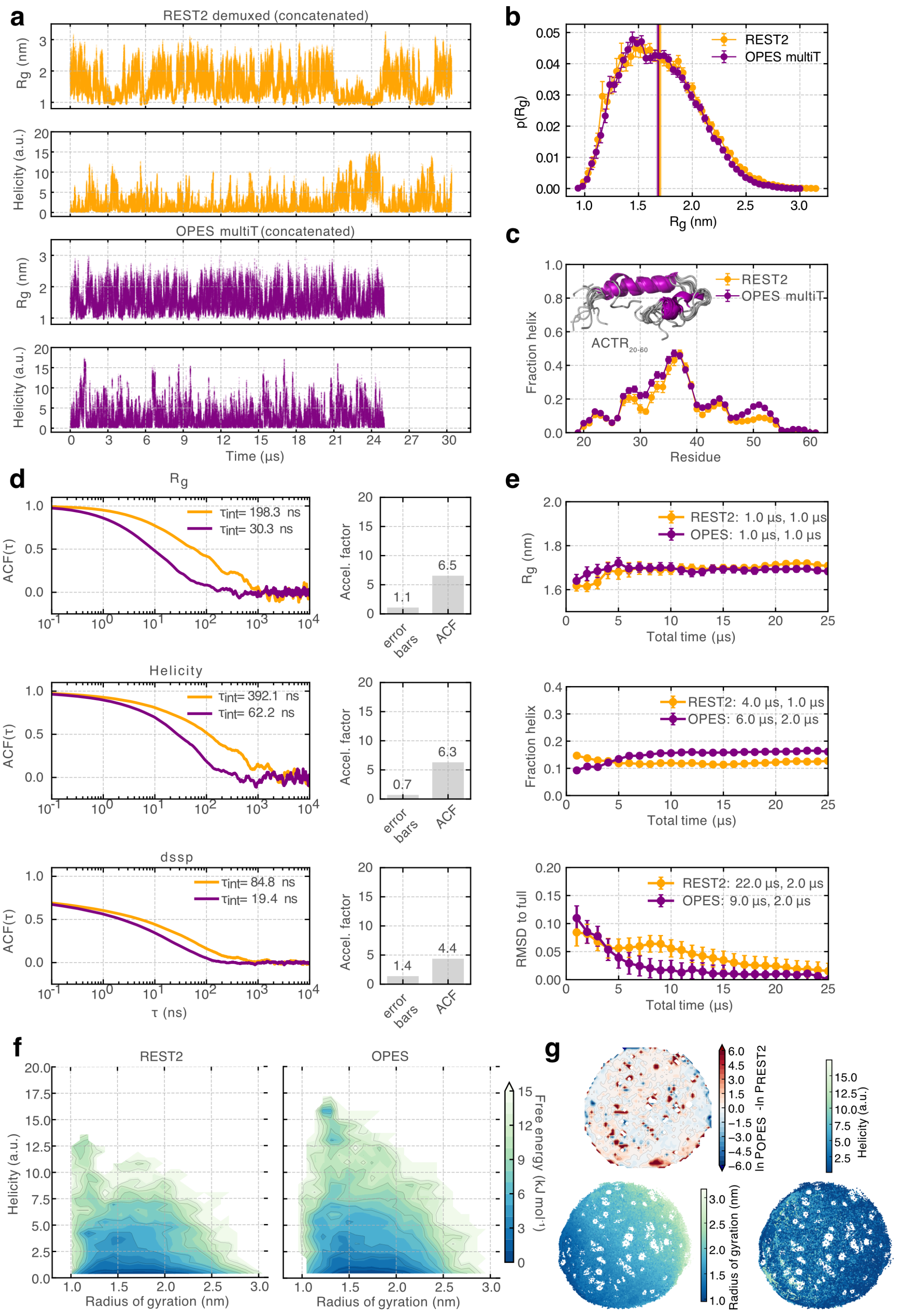
a
REST2 demuxed (concatenated)
Rg (nm)
Helicity (a.u.)
OPES multiT (concatenated)
Time (µs)
b
REST2
OPES multiT
p(Rg)
Rg (nm)
c
REST2
OPES multiT
ACTR20-60
Fraction helix
Residue
d
Rg
τint= 198.3 ns
τint= 30.3 ns
ACF(τ)
Accel. factor
error bars
ACF
Helicity
τint= 392.1 ns
τint= 62.2 ns
dssp
τint= 84.8 ns
τint= 19.4 ns
τ (ns)
e
REST2: 1.0 µs, 1.0 µs
OPES: 1.0 µs, 1.0 µs
Total time (µs)
REST2: 4.0 µs, 1.0 µs
OPES: 6.0 µs, 2.0 µs
REST2: 22.0 µs, 2.0 µs
OPES: 9.0 µs, 2.0 µs
RMSD to full
f
REST2
OPES
Helicity (a.u.)
Radius of gyration (nm)
Free energy (kJ mol⁻¹)
g
ln POPES -ln PREST2
Helicity (a.u.)
Radius of gyration (nm)

**Figure S11.** OPES multiT and REST2 simulations of ACTR$_{20-60}$. **a.** Fluctuations of the radius of gyration ($R_g$) and protein helicity sampled with REST2 and OPES multithermal simulations. **b.** Probability distributions and ensemble-averaged values (vertical lines) of the $R_g$. Errors represent the standard error of the mean calculated with a blocking analysis **c.** Mean and standard errors of per-residue helical fractions. **d.** Normalised autocorrelation function (ACF) of the $R_g$ (top), helicity (bottom) and per-residue secondary structure (dssp) calculated from the demuxed REST2 and OPES multiT timeseries sampled every 40 ps. The correlation times were estimated using the integral of the ACF ($\tau_{int}$). The right panels show estimates of an acceleration factor (OPES / REST2) estimated from the error bars of ensemble-averaged values and correlation times (ACF). **e.** Time-cumulative averages of the $R_g$, total fraction helix and per-residue helical fractions (measuring the RMSD to the profile derived from the full ensemble) are shown together with the extracted stabilisation times, which mark the earliest points at which the running averages and their uncertainties, respectively, converge to stable values within predefined tolerances. **f.** Free-energy landscapes of ensembles sampled at 300 K by both methods as a function of $R_g$ and helicity. **g.** 2D ELViM projections of a combined REST2 + OPES ensemble at 300 K (see Methods). Top: difference in log probabilities across the entire landscape. Bottom: projections coloured by $R_g$ and helicity.

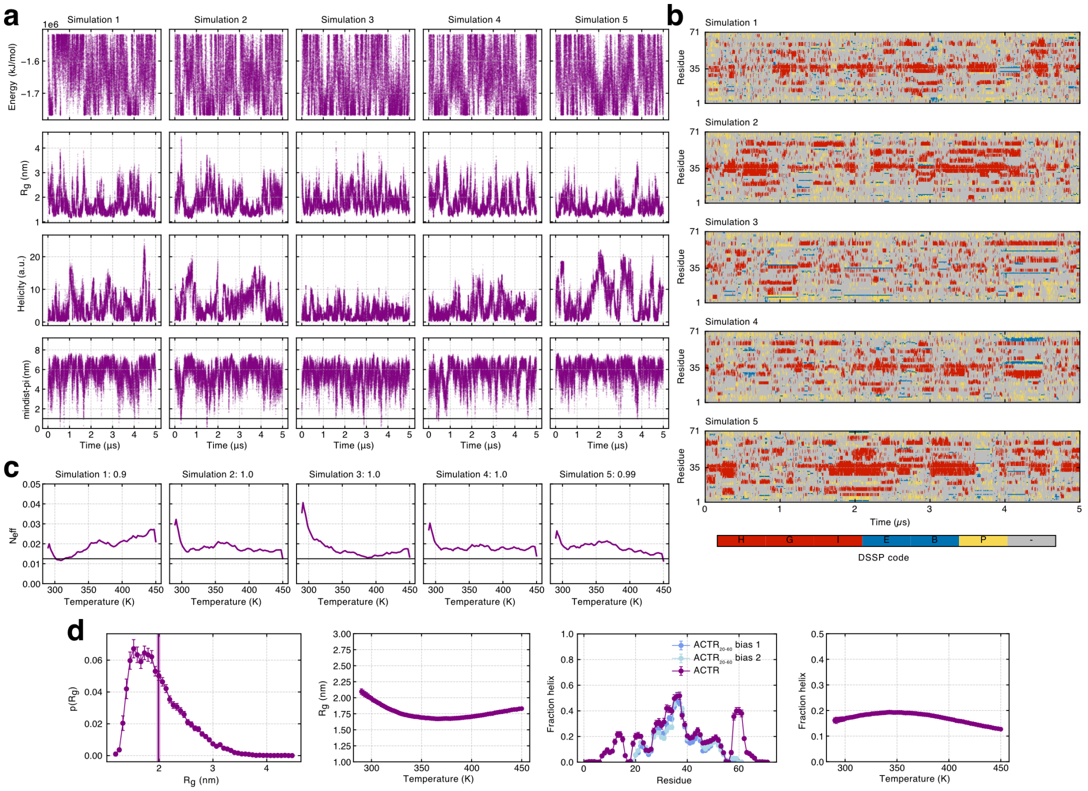


**Figure S12.** OPES simulations of ACTR. **a.** Fluctuations of the system potential energy, protein radius of gyration ($R_g$), total helicity, and minimum distance to a periodic image sampled with OPES multithermal simulations and one shared, static bias potential. **b.** Time-resolved secondary structure of the protein coloured according to per-residue DSSP assignment. **c.** Effective sample sizes across the range of temperatures sampled in the simulations and sampling scores from panels a-b. **d.** Probability distribution of the $R_g$ at 300 K, average $R_g$ values as function of temperature, per-residue helical fractions at 300 K, and total helical fraction as a function of temperature. Per-residue helical fraction of ACTR$_{20-60}$ obtained from OPES multithermal simulations with two different biases are overlaid with full-length ACTR for a comparison.

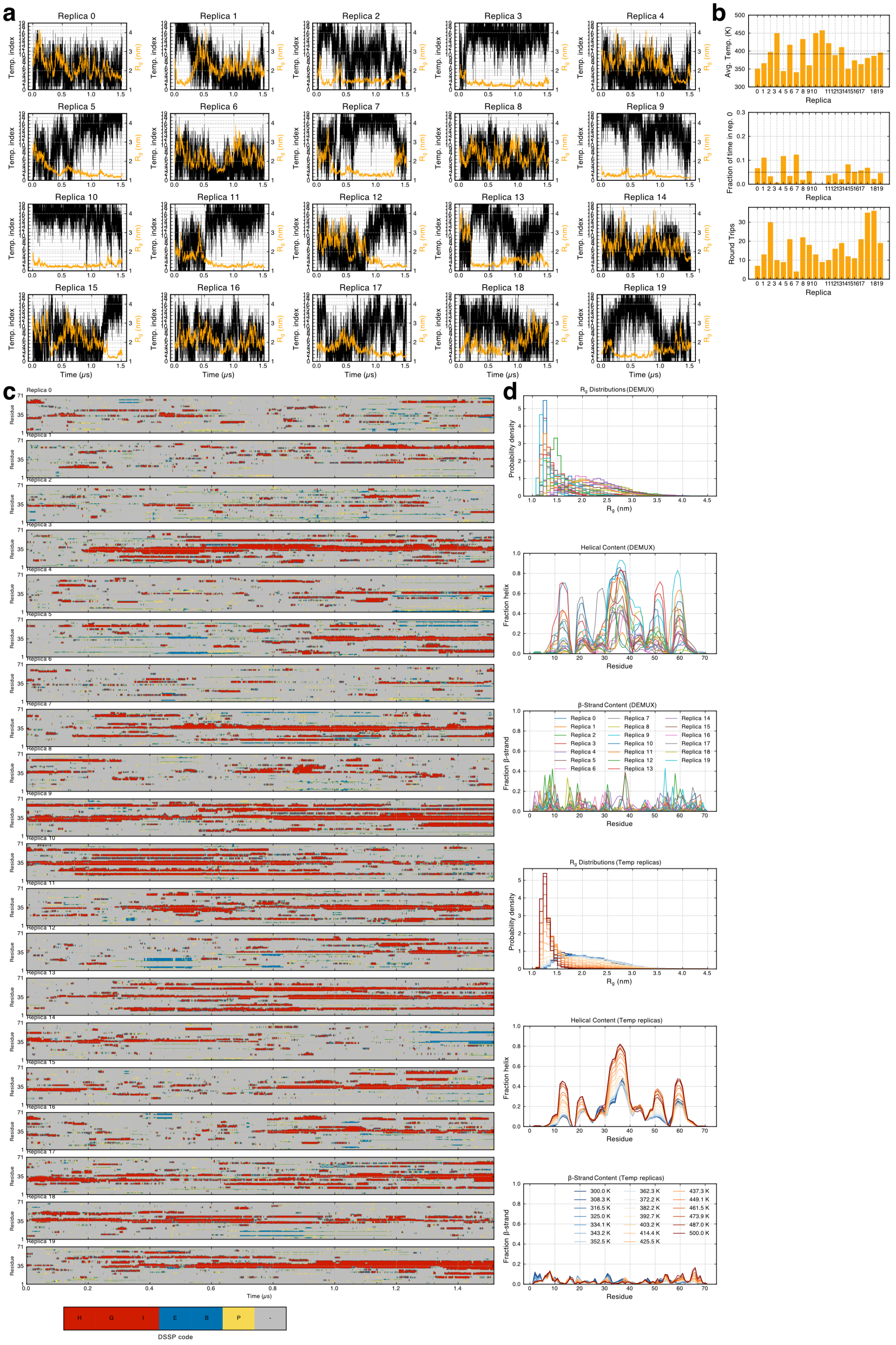

**Figure S13.** REST2 simulations of ACTR. **a.** Demuxed time series of all replicas showing their traversal through temperature replicas (left axis, black) and fluctuations of the $R_g$ (right axis, orange). **b.** Assessment of REST2 replica mixing. Top: average temperature of each demuxed replica over the course of the entire simulation. Middle: fraction of time spent in the lowest temperature replica. Bottom: number of round trips through temperature space. **c.** Time-resolved secondary structure of the protein calculated from demuxed trajectories and coloured according to per-residue DSSP assignment. **d.** $R_g$ distributions, per-residue helical fractions and per-residue $\beta$-strand fractions (top to bottom, respectively) calculated from demuxed and temperature replicas.

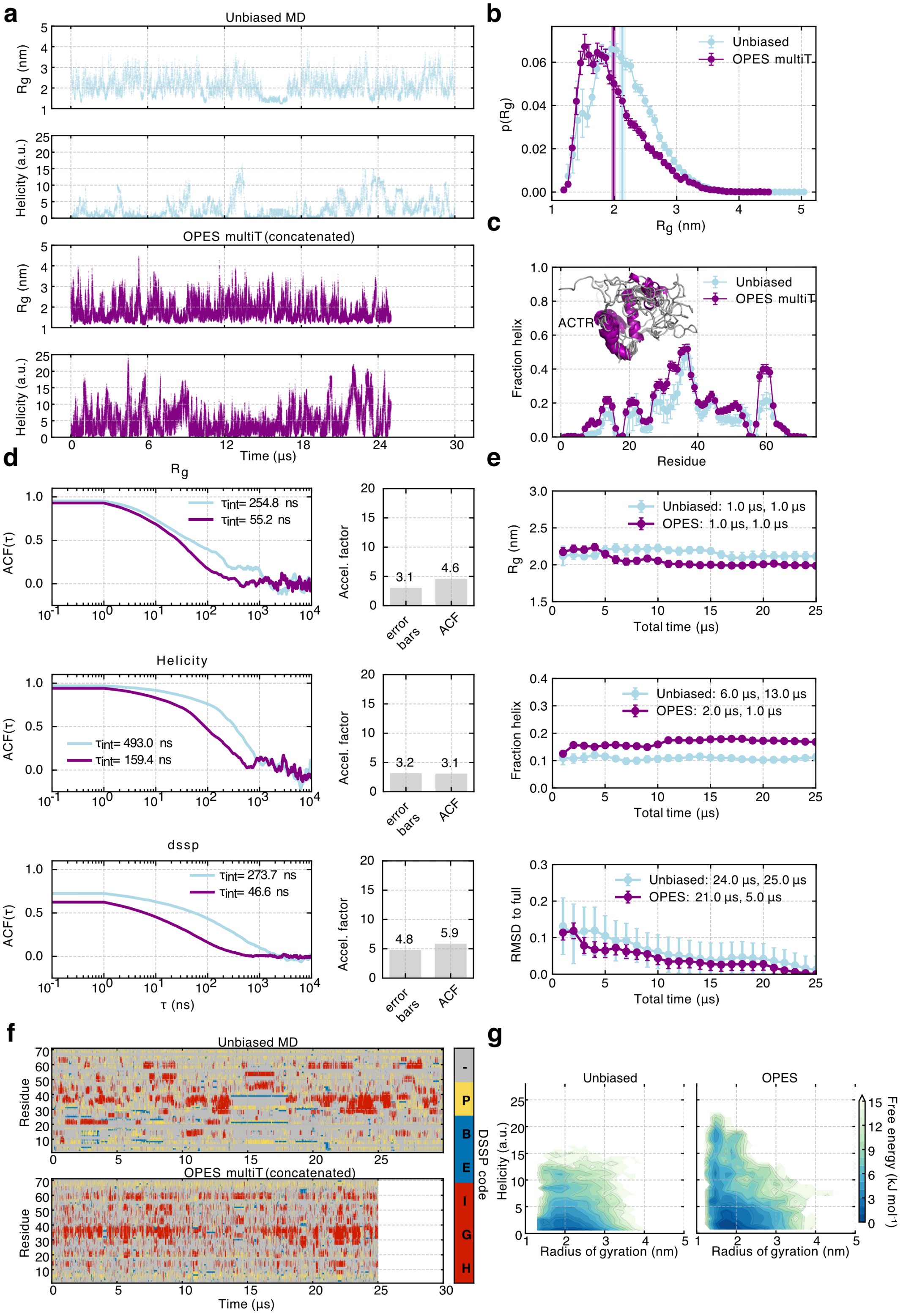

a
Unbiased MD
OPES multiT (concatenated)
Time (μs)
b
Unbiased
OPES multiT
c
ACTR
Fraction helix
Residue
d
Rg
Helicity
dssp
ACF(τ)
τ (ns)
Accel. factor
error bars
ACF
e
Unbiased: 1.0 μs, 1.0 μs
OPES: 1.0 μs, 1.0 μs
Unbiased: 6.0 μs, 13.0 μs
OPES: 2.0 μs, 1.0 μs
Unbiased: 24.0 μs, 25.0 μs
OPES: 21.0 μs, 5.0 μs
Total time (μs)
RMSD to full
f
Residue
DSSP code
g
Unbiased
OPES
Radius of gyration (nm)
Helicity (a.u.)
Free energy (kJ mol⁻¹)

**Figure S14.** Comparison with unbiased simulations of ACTR. **a.** Fluctuations of the radius of gyration ($R_g$) and protein helicity sampled with unbiased (***Robustelli et al., 2018***) and OPES multithermal simulations. **b.** Probability distributions and ensemble-averaged values (vertical lines) of the $R_g$. Errors represent the standard error of the mean calculated with a blocking analysis **c.** Mean and standard errors of per-residue helical fractions. **d.** Normalised autocorrelation function (ACF) of the $R_g$ (top), helicity (bottom) and per-residue secondary structure (dssp) calculated from the unbiased and OPES multiT timeseries sampled every 1 ns. The correlation times were estimated using the integral of the ACF ($\tau_{int}$). The right panels show estimates of an acceleration factor (OPES / unbiased) estimated from the error bars of ensemble-averaged values and correlation times (ACF). **e.** Time-cumulative averages of the $R_g$, total fraction helix and per-residue helical fractions (measuring the RMSD to the profile derived from the full ensemble) are shown together with the extracted stabilisation times, which mark the earliest points at which the running averages and their uncertainties, respectively, converge to stable values within predefined tolerances. **f.** Time-resolved secondary structure of the protein coloured according to per-residue DSSP assignment. **g.** Free-energy landscapes of ensembles sampled at 300 K by both methods as a function of $R_g$ and helicity.

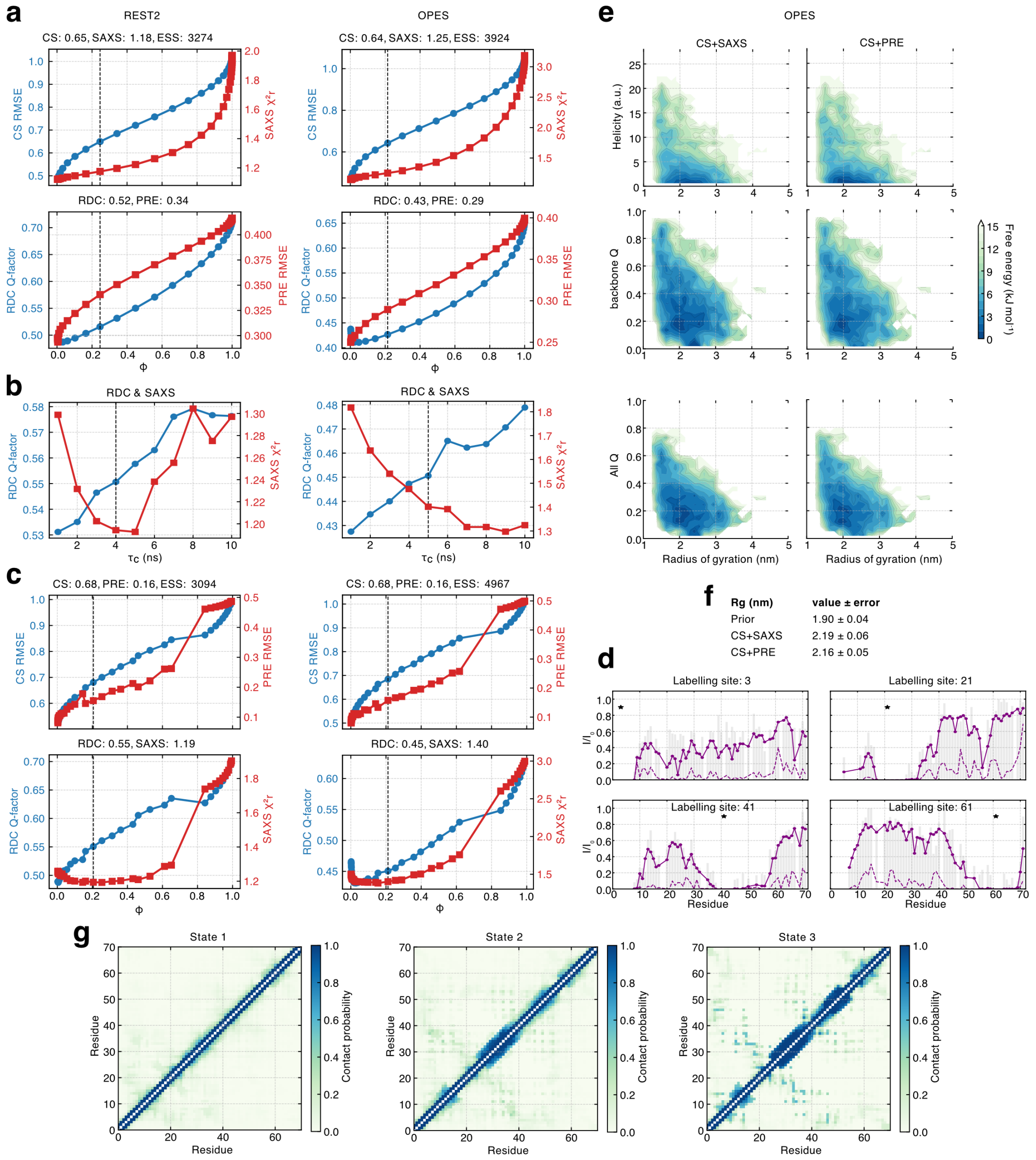

**Figure S15.** Ensemble reweighting of ACTR with experimental data. **a.** Reweighting curves showing agreement with active data (top panel, CS+SAXS) and passive data (bottom panel, RDC+PRE) for the REST2 and OPES ensembles as a function of $\phi$. The agreement with all data types (CS: normalised RMSE, SAXS: reduced $\chi^2$, RDC: Q-factor, PRE: RMSE) and effective sample size (ESS) after reweighting for the indicated posterior (vertical lines) are annotated above each plot. **b.** Agreement with passive data (RDC+SAXS) after reweighting with CS+PRE using different values of the rotational correlation time, $\tau_c$. A suitable value was selected on the basis of resulting in a compromise between the RDC and SAXS data (4 and 5 ns for REST2 and OPES ensembles, respectively). **c.** Reweighting curves showing agreement with active data (top panel, CS+PRE) and passive data (bottom panel, RDC+SAXS) for the REST2 and OPES ensembles. **d.** Agreement between experimental (grey bars) and calculated PRE intensity ratios for all labelling sites plotted across the sequence for the reweighted OPES ensemble from panel c. **e.** Free-energy landscapes of OPES ensembles after reweighting with CS+SAXS (left) and CS+PRE (right) data as a function of helicity (top), backbone native contacts (middle) and all native contacts (bottom) of the bound state (PDB 1KBH) and $R_g$ at 310 K. **f.** Ensemble-averaged value (mean ± s.e.m.) of the $R_g$ before and after reweighting (OPES ensemble). **g.** Intramolecular contact probability maps of the main conformational states (low energy basins) of the reweighted ACTR ensemble (OPES) using a heavy atom distance cut-off of 0.6 nm.

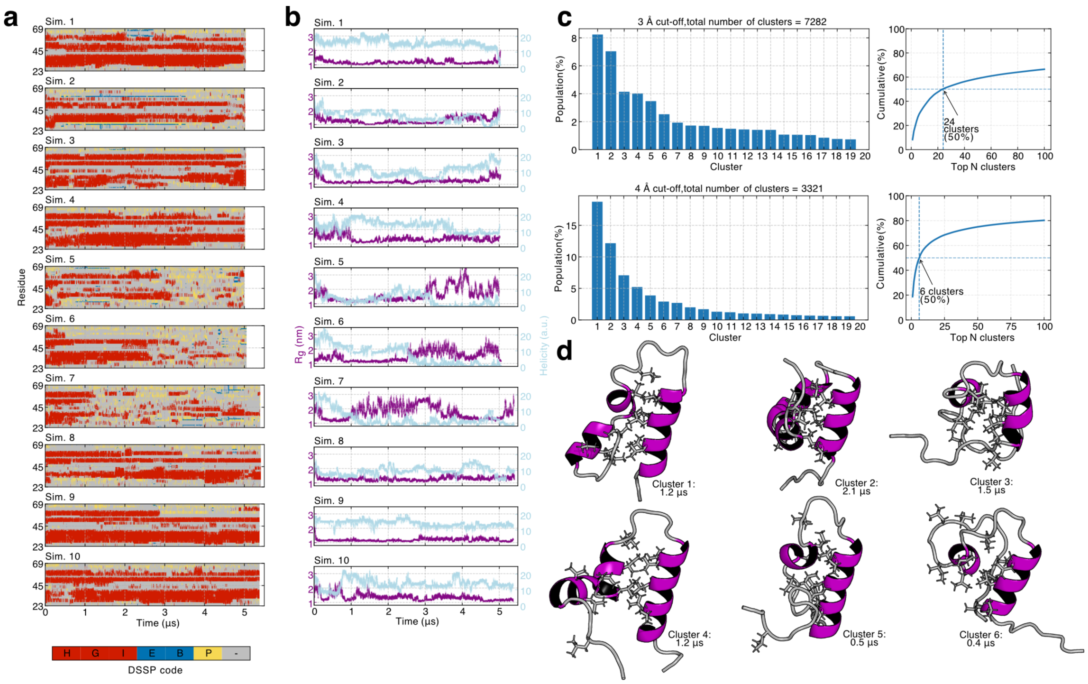


**Figure S16.** Unbiased MD simulations of ACTR (residues 23-69) initiated from the experimental structure of the bound state (PDB 1KBH, without CBP). **a.** Time-resolved secondary structure of the protein coloured according to per-residue DSSP assignment. **b.** Fluctuations of the $R_g$ and protein helicity. **c.** Conformational cluster populations obtained with two different cut-off values indicated above each plot using backbone atoms of residues 27–63 for alignment and RMSD calculations (see Methods). The right plot shows the cumulative population captured by the top N clusters and the number of clusters required to capture 50% of the ensemble. **d.** Structures of the top 6 cluster centroids with a 4 Å cut-off. The maximum observed lifetimes are indicated for each cluster using the cluster-frame assignments (continuous time segments were defined with a time gap allowance of 20 ns).

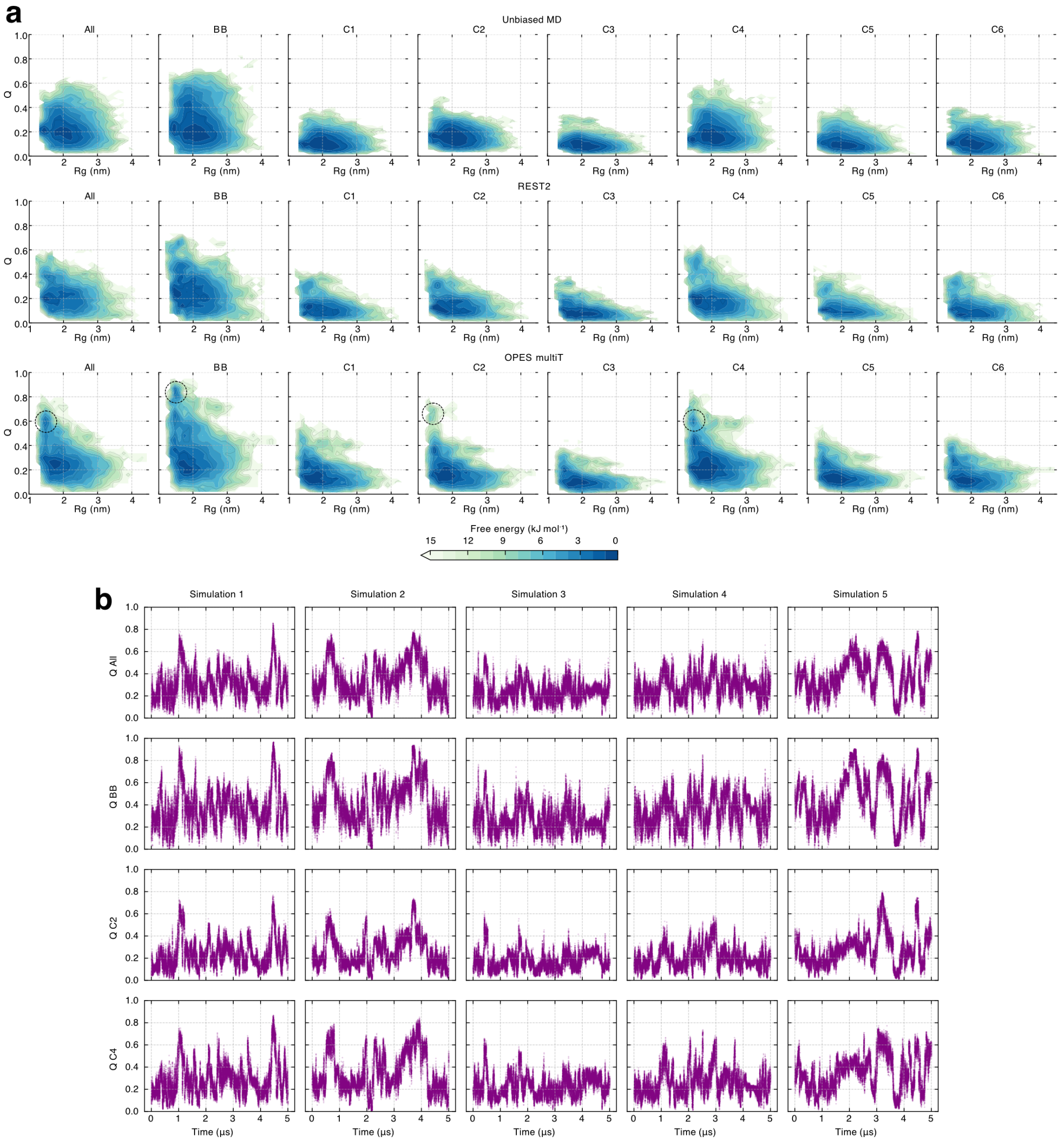


**Figure S17.** Sampling of native contacts from different reference structures in unbiased, REST2 and OPES multithermal simulations of ACTR. **a.** Free-energy landscapes at 300 K as a function of $R_g$ and Q. All and BB refer to all heavy atom and backbone native contacts of the bound state, respectively (PDB 1KBH). C1–6 refer to cluster centroids of obtained from unbiased MD simulations presented in Figure S16. Circles indicate local free-energy minima sampled by OPES not explored by the other methods for All, BB, C2, and C4. **b.** Fluctuations of All, BB, C2, and C4 Q values as a function of time in all OPES multithermal replicas of ACTR, highlighting multiple transitions between low and high Q regions.

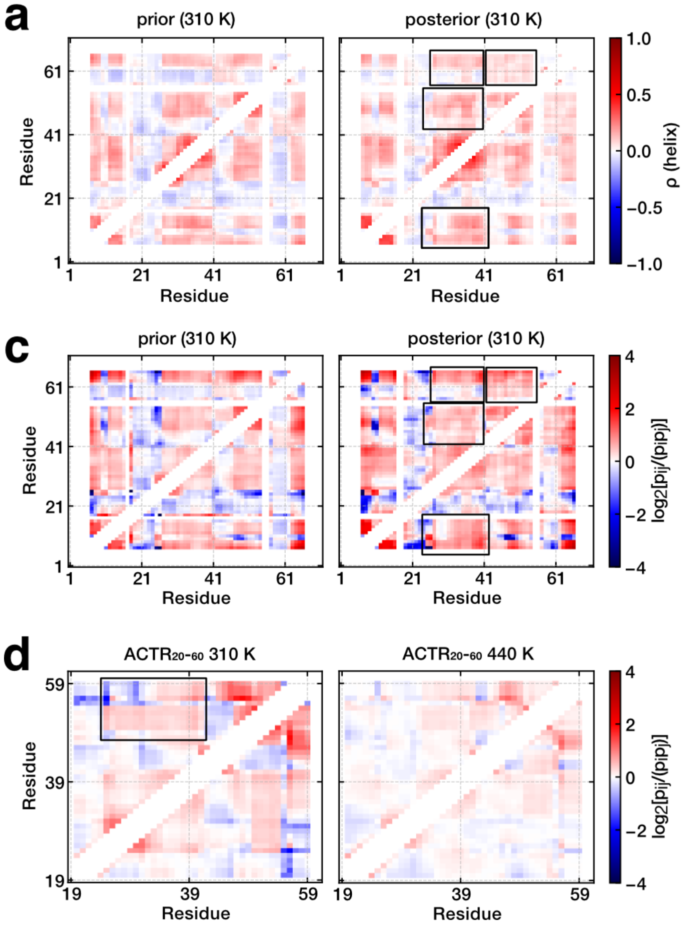


b

| Threshold | 0.5 | 0.6 | 0.7 | 0.8 |
|---|---|---|---|---|
| **Helices** | | | | |
| 0-1 | 2.25 | 3.78 | nan | nan |
| | *0.030* | *0.005* | *0.000* | *0.000* |
| 0-2 | 1.72 | 2.63 | nan | nan |
| | *0.007* | *0.001* | *0.000* | *0.000* |
| 0-3 | 0.37 | 0.50 | nan | nan |
| | *0.001* | *0.000* | *0.000* | *0.000* |
| 1-2 | 2.40 | 2.91 | 3.53 | 4.42 |
| | *0.061* | *0.037* | *0.021* | *0.008* |
| 1-3 | 1.17 | 1.17 | 2.00 | 1.74 |
| | *0.017* | *0.013* | *0.005* | *0.001* |
| 2-3 | 2.03 | 2.78 | 2.67 | 1.25 |
| | *0.009* | *0.008* | *0.002* | *0.000* |
| 1-2-3 | 6.42 | 10.34 | 16.66 | 1.33 |
| | *0.008* | *0.006* | *0.002* | *0.000* |
| **p(fold)** | | | | |
| **Helix** | | | | |
| 0 | *0.047* | *0.007* | *0.000* | *0.000* |
| 1 | *0.282* | *0.210* | *0.148* | *0.119* |
| 2 | *0.090* | *0.055* | *0.039* | *0.015* |
| 3 | *0.052* | *0.051* | *0.016* | *0.005* |

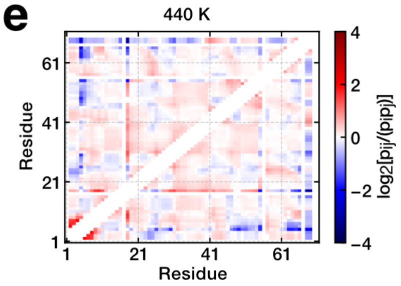


**Figure S18.** Cooperativity of helix formation in ACTR. **a.** Pearson correlation coefficient ($\rho$) matrix of helix formation calculated with DSSP (0 = no helix, 1 = helix) before and after ensemble reweighting with experimental data. Boxes highlight regions corresponding to long-range, correlated folding of helices 0 (residues 6–18), 1 (residues 27–38), 2 (residues 47–55), and 3 (residues 57–64), defined based on regions exhibiting secondary structure by NMR (***Iešmantavičius et al., 2013***). Only residues separated by at least 4 positions and having a helical probability of at least 0.01 were included. **b.** Cooperativity ( $p_{ij}$ / $(p_i p_j)$ ) of helix formation of different pairs/groups of the same helices (0–3) after ensemble reweighting. Different thresholds defining complete helix formation (fraction of residues simultaneously in a helical state within a given helical region) were used. Values shown in *italics* represent the observed joint probabilities (i.e., multiple helices formed simultaneously). The bottom table shows the total folding probabilities of the individual helices with the different thresholds. **c.** Cooperativity ( $p_{ij}$ / $(p_i p_j)$ ) matrix of helix formation before and after reweighting. Boxes highlight regions corresponding to long-range, correlated folding of helices 0 (residues 6–18), 1 (residues 27–38), 2 (residues 47–55), and 3 (residues 57–64). Only residues separated by at least 4 positions and having a helical probability of at least 0.01 were included. **d.** Cooperativity matrix of $ACTR_{20-60}$ OPES simulations reweighted (with the OPES bias) at 310 and 440 K. The box indicates the cooperative formation of helices 1 and 2. **e.** Cooperativity matrix of full-length ACTR reweighted (with the OPES bias) at 440 K. The reduction in cooperativities at 440 relative to 310 K is indicative of a loss in long-range tertiary interactions and folded structure.

#### Supplementary tables

| System | unbiased MD | OPES | decrease (%) |
|---|---|---|---|
| $(AAQAA)_3$ | 312.0 ± 1.6 | 193.3 ± 0.6 | 38.0 ± 0.2 |
| $ACTR_{20-60}$ | 199.0 ± 1.5 | 136.7 ± 0.4 | 31.3 ± 0.2 |
| ACTR | 108.0 ± 0.4 | 77.2 ± 0.3 | 28.5 ± 0.1 |
| HTTex1 16Q | 63.9 ± 0.3 | 44.8 ± 0.2 | 29.9 ± 0.2 |

**Table S1.** Comparison of unbiased MD and OPES simulation performances (mean ± std. dev. measured from 3 simulations in ns/day using one V100 GPU).

| System | N | trial range (min-max) | $N_{TS}$ | success rate (%) |
|---|---|---|---|---|
| $(AAQAA)_3$ 269-450 K | 29,730 | 57–73 | 55 | 30 |
| $(AAQAA)_3$ 290-450 K | 29,730 | 52–75 | 48 | 50 |
| $ACTR_{20-60}$ 290-450 K | 47,040 | 67–80 | 70 | 20 |
| ACTR 290-450 K | 95,440 | 87–116 | 80 | 10 |
| HTTex1 16Q 288-450 K | 174,602 | 118–152 | 100 | 20 |

**Table S2.** Summary of simulation systems, number of atoms (N), range of the number of temperature steps ($N_{TS}$) from ten automatic trial simulations (where the number of temperature steps is determined automatically by PLUMED based on the potential energy fluctuations (***Invernizzi et al., 2020***)), the used number of temperature steps for production simulations ($N_{TS}$), and success rate in obtaining a converged bias potential out of ten trial simulations with a fixed number of $N_{TS}$. A linear regression of $\sqrt{N}$ versus $N_{TS}$ yields a slope of 0.192 and an intercept of 20.8 (Pearson r = 0.966). The required number of temperature steps for the temperature range 290–450 K can therefore be estimated as $0.192\sqrt{N} + 20.8$.

| Bins | Eff. coverage OPES | Eff. coverage REST2 | ΔEff. coverage (%) | Norm. entropy OPES | Norm. entropy REST2 | ΔNorm. entropy (%) | Rel. coverage OPES\|REST2 | Rel. coverage REST2\|OPES | ΔRel. coverage (%) |
|---|---|---|---|---|---|---|---|---|---|
| 21 | 0.4046 ± 0.0046 | 0.3913 ± 0.0045 | -3.39 ± 1.66 | 0.8998 ± 0.0010 | 0.8969 ± 0.0011 | -0.32 ± 0.17 | 0.9851 ± 0.0048 | 0.9758 ± 0.0050 | -0.95 ± 0.71 |
| 43 | 0.3158 ± 0.0040 | 0.3121 ± 0.0038 | -1.18 ± 1.78 | 0.8925 ± 0.0009 | 0.8919 ± 0.0009 | -0.07 ± 0.15 | 0.9238 ± 0.0054 | 0.9301 ± 0.0049 | 0.68 ± 0.78 |
| 65 | 0.2713 ± 0.0034 | 0.2727 ± 0.0034 | 0.52 ± 1.78 | 0.8848 ± 0.0008 | 0.8862 ± 0.0009 | 0.16 ± 0.14 | 0.8440 ± 0.0048 | 0.8627 ± 0.0050 | 2.17 ± 0.80 |
| 87 | 0.2422 ± 0.0029 | 0.2439 ± 0.0029 | 0.72 ± 1.67 | 0.8771 ± 0.0008 | 0.8794 ± 0.0008 | 0.26 ± 0.12 | 0.7614 ± 0.0048 | 0.7849 ± 0.0048 | 2.99 ± 0.86 |
| 108 | 0.2129 ± 0.0024 | 0.2206 ± 0.0026 | 3.48 ± 1.60 | 0.8668 ± 0.0007 | 0.8714 ± 0.0007 | 0.52 ± 0.12 | 0.6813 ± 0.0046 | 0.7118 ± 0.0049 | 4.29 ± 0.94 |
| 130 | 0.1858 ± 0.0021 | 0.1963 ± 0.0022 | 5.38 ± 1.55 | 0.8559 ± 0.0007 | 0.8620 ± 0.0007 | 0.71 ± 0.12 | 0.5986 ± 0.0045 | 0.6332 ± 0.0045 | 5.47 ± 1.00 |
| 152 | 0.1641 ± 0.0017 | 0.1744 ± 0.0018 | 5.92 ± 1.43 | 0.8459 ± 0.0007 | 0.8524 ± 0.0007 | 0.77 ± 0.11 | 0.5299 ± 0.0040 | 0.5633 ± 0.0045 | 5.92 ± 1.07 |

**Table S3.** Quantification of conformational space exploration of REST2 and OPES simulations of HTTex1 16Q using 2D ELViM projections. Errors represent standard deviations obtained with bootstrapping.

| Bins | Eff. coverage OPES | Eff. coverage REST2 | ΔEff. coverage (%) | Norm. entropy OPES | Norm. entropy REST2 | ΔNorm. entropy (%) | Rel. coverage OPES\|REST2 | Rel. coverage REST2\|OPES | ΔRel. coverage (%) |
|---|---|---|---|---|---|---|---|---|---|
| 21 | 0.1337 ± 0.0019 | 0.1059 ± 0.0014 | -26.29 ± 2.24 | 0.7726 ± 0.0016 | 0.7106 ± 0.0015 | -8.74 ± 0.31 | 0.9862 ± 0.0055 | 0.6363 ± 0.0093 | -54.98 ± 1.88 |
| 43 | 0.0958 ± 0.0016 | 0.0751 ± 0.0012 | -27.59 ± 2.66 | 0.7936 ± 0.0014 | 0.7429 ± 0.0014 | -6.84 ± 0.26 | 0.9593 ± 0.0057 | 0.6458 ± 0.0069 | -48.54 ± 1.49 |
| 65 | 0.0819 ± 0.0014 | 0.0637 ± 0.0011 | -28.60 ± 2.85 | 0.7997 ± 0.0013 | 0.7568 ± 0.0013 | -5.66 ± 0.24 | 0.9206 ± 0.0055 | 0.6460 ± 0.0059 | -42.49 ± 1.31 |
| 87 | 0.0734 ± 0.0013 | 0.0566 ± 0.0010 | -29.66 ± 3.02 | 0.8003 ± 0.0012 | 0.7631 ± 0.0012 | -4.88 ± 0.22 | 0.8637 ± 0.0057 | 0.6333 ± 0.0054 | -36.37 ± 1.28 |
| 108 | 0.0672 ± 0.0012 | 0.0515 ± 0.0009 | -30.48 ± 3.09 | 0.7974 ± 0.0011 | 0.7650 ± 0.0012 | -4.24 ± 0.21 | 0.7932 ± 0.0059 | 0.6061 ± 0.0051 | -30.87 ± 1.32 |
| 130 | 0.0628 ± 0.0012 | 0.0469 ± 0.0009 | -33.86 ± 3.22 | 0.7929 ± 0.0011 | 0.7648 ± 0.0011 | -3.68 ± 0.20 | 0.7056 ± 0.0056 | 0.5674 ± 0.0048 | -24.36 ± 1.33 |
| 152 | 0.0583 ± 0.0011 | 0.0436 ± 0.0008 | -33.77 ± 3.22 | 0.7876 ± 0.0010 | 0.7633 ± 0.0011 | -3.19 ± 0.19 | 0.6305 ± 0.0055 | 0.5264 ± 0.0047 | -19.78 ± 1.38 |

**Table S4.** Quanitification of conformational space exploration of REST2 and OPES simulations of HTTex1 16Q using the $R_g$ and helicity CVs for 2D projections. Errors represent standard deviations obtained with bootstrapping.

| Bins | Eff. coverage OPES | Eff. coverage REST2 | ΔEff. coverage (%) | Norm. entropy OPES | Norm. entropy REST2 | ΔNorm. entropy (%) | Rel. coverage OPES\|REST2 | Rel. coverage REST2\|OPES | ΔRel. coverage (%) |
|---|---|---|---|---|---|---|---|---|---|
| 27 | 0.3969 ± 0.0036 | 0.4008 ± 0.0034 | 0.98 ± 1.23 | 0.9026 ± 0.0007 | 0.9034 ± 0.0007 | 0.09 ± 0.11 | 0.9785 ± 0.0040 | 0.9847 ± 0.0036 | 0.63 ± 0.55 |
| 55 | 0.3282 ± 0.0033 | 0.3335 ± 0.0031 | 1.60 ± 1.35 | 0.9006 ± 0.0006 | 0.9017 ± 0.0006 | 0.12 ± 0.10 | 0.9216 ± 0.0041 | 0.9334 ± 0.0036 | 1.27 ± 0.59 |
| 83 | 0.2909 ± 0.0030 | 0.2983 ± 0.0026 | 2.48 ± 1.31 | 0.8962 ± 0.0006 | 0.8976 ± 0.0006 | 0.15 ± 0.09 | 0.8567 ± 0.0039 | 0.8704 ± 0.0039 | 1.58 ± 0.63 |
| 111 | 0.2577 ± 0.0025 | 0.2690 ± 0.0024 | 4.22 ± 1.30 | 0.8887 ± 0.0006 | 0.8914 ± 0.0006 | 0.30 ± 0.09 | 0.7765 ± 0.0038 | 0.7938 ± 0.0038 | 2.18 ± 0.67 |
| 138 | 0.2298 ± 0.0022 | 0.2411 ± 0.0021 | 4.66 ± 1.24 | 0.8800 ± 0.0005 | 0.8834 ± 0.0005 | 0.39 ± 0.08 | 0.7009 ± 0.0036 | 0.7221 ± 0.0036 | 2.94 ± 0.71 |
| 166 | 0.2021 ± 0.0018 | 0.2134 ± 0.0016 | 5.29 ± 1.15 | 0.8700 ± 0.0005 | 0.8742 ± 0.0005 | 0.48 ± 0.08 | 0.6224 ± 0.0034 | 0.6459 ± 0.0035 | 3.64 ± 0.75 |
| 194 | 0.1774 ± 0.0014 | 0.1892 ± 0.0015 | 6.27 ± 1.08 | 0.8597 ± 0.0005 | 0.8646 ± 0.0005 | 0.57 ± 0.08 | 0.5539 ± 0.0030 | 0.5779 ± 0.0035 | 4.15 ± 0.80 |

**Table S5.** Quanitification of conformational space exploration of REST2 and OPES simulations of the $ACTR_{20-60}$ fragment using 2D ELViM projections. Errors represent standard deviations obtained with bootstrapping.

| Bins | Eff. coverage OPES | Eff. coverage REST2 | ΔEff. coverage (%) | Norm. entropy OPES | Norm. entropy REST2 | ΔNorm. entropy (%) | Rel. coverage OPES\|REST2 | Rel. coverage REST2\|OPES | ΔRel. coverage (%) |
|---|---|---|---|---|---|---|---|---|---|
| 27 | 0.1425 ± 0.0015 | 0.0997 ± 0.0009 | -42.93 ± 1.85 | 0.7865 ± 0.0010 | 0.7255 ± 0.0011 | -8.41 ± 0.21 | 0.9622 ± 0.0068 | 0.7621 ± 0.0089 | -26.27 ± 1.50 |
| 55 | 0.1295 ± 0.0015 | 0.0886 ± 0.0010 | -46.23 ± 2.08 | 0.8155 ± 0.0009 | 0.7650 ± 0.0009 | -6.61 ± 0.16 | 0.9177 ± 0.0055 | 0.6882 ± 0.0061 | -33.34 ± 1.24 |
| 83 | 0.1233 ± 0.0015 | 0.0837 ± 0.0009 | -47.44 ± 2.16 | 0.8245 ± 0.0008 | 0.7800 ± 0.0008 | -5.70 ± 0.14 | 0.8763 ± 0.0048 | 0.6584 ± 0.0048 | -33.08 ± 1.05 |
| 111 | 0.1175 ± 0.0014 | 0.0814 ± 0.0009 | -44.29 ± 2.06 | 0.8263 ± 0.0007 | 0.7878 ± 0.0007 | -4.88 ± 0.13 | 0.8341 ± 0.0042 | 0.6394 ± 0.0041 | -30.46 ± 0.94 |
| 138 | 0.1114 ± 0.0012 | 0.0788 ± 0.0009 | -41.46 ± 1.96 | 0.8250 ± 0.0006 | 0.7906 ± 0.0007 | -4.35 ± 0.12 | 0.7928 ± 0.0042 | 0.6158 ± 0.0038 | -28.74 ± 0.94 |
| 166 | 0.1041 ± 0.0011 | 0.0751 ± 0.0008 | -38.57 ± 1.89 | 0.8210 ± 0.0006 | 0.7913 ± 0.0007 | -3.75 ± 0.11 | 0.7387 ± 0.0041 | 0.5897 ± 0.0037 | -25.25 ± 0.95 |
| 194 | 0.0967 ± 0.0010 | 0.0701 ± 0.0008 | -37.98 ± 1.87 | 0.8164 ± 0.0006 | 0.7897 ± 0.0006 | -3.38 ± 0.11 | 0.6902 ± 0.0040 | 0.5617 ± 0.0035 | -22.88 ± 0.96 |

**Table S6.** Quanitification of conformational space exploration of REST2 and OPES simulations of the $ACTR_{20-60}$ fragment using the $R_g$ and helicity CVs for 2D projections. Errors represent standard deviations obtained with bootstrapping.

| Bins | Eff. coverage OPES | Eff. coverage REST2 | ΔEff. coverage (%) | Norm. entropy OPES | Norm. entropy REST2 | ΔNorm. entropy (%) | Rel. coverage OPES\|REST2 | Rel. coverage REST2\|OPES | ΔRel. coverage (%) |
|---|---|---|---|---|---|---|---|---|---|
| 25 | 0.3667 ± 0.0040 | 0.2812 ± 0.0032 | -30.40 ± 1.85 | 0.8959 ± 0.0009 | 0.8713 ± 0.0010 | -2.82 ± 0.15 | 0.9700 ± 0.0041 | 0.9502 ± 0.0052 | -2.08 ± 0.70 |
| 50 | 0.2814 ± 0.0033 | 0.2200 ± 0.0026 | -27.90 ± 1.94 | 0.8879 ± 0.0008 | 0.8667 ± 0.0009 | -2.44 ± 0.13 | 0.8940 ± 0.0047 | 0.8455 ± 0.0048 | -5.74 ± 0.80 |
| 75 | 0.2423 ± 0.0028 | 0.1908 ± 0.0023 | -27.03 ± 1.92 | 0.8808 ± 0.0007 | 0.8628 ± 0.0008 | -2.09 ± 0.12 | 0.8081 ± 0.0047 | 0.7582 ± 0.0046 | -6.58 ± 0.87 |
| 101 | 0.2121 ± 0.0025 | 0.1737 ± 0.0021 | -22.09 ± 1.90 | 0.8728 ± 0.0007 | 0.8579 ± 0.0007 | -1.74 ± 0.12 | 0.7174 ± 0.0043 | 0.6748 ± 0.0039 | -6.32 ± 0.86 |
| 126 | 0.1916 ± 0.0021 | 0.1586 ± 0.0018 | -20.79 ± 1.77 | 0.8650 ± 0.0006 | 0.8519 ± 0.0007 | -1.53 ± 0.11 | 0.6334 ± 0.0042 | 0.5969 ± 0.0037 | -6.12 ± 0.94 |
| 151 | 0.1708 ± 0.0018 | 0.1448 ± 0.0016 | -17.98 ± 1.65 | 0.8558 ± 0.0006 | 0.8452 ± 0.0006 | -1.25 ± 0.10 | 0.5604 ± 0.0040 | 0.5342 ± 0.0036 | -4.92 ± 1.02 |
| 176 | 0.1525 ± 0.0016 | 0.1321 ± 0.0014 | -15.47 ± 1.60 | 0.8471 ± 0.0006 | 0.8379 ± 0.0006 | -1.10 ± 0.10 | 0.4991 ± 0.0039 | 0.4760 ± 0.0035 | -4.85 ± 1.10 |

**Table S7.** Quanitification of conformational space exploration of REST2 and OPES simulations of ACTR using 2D ELViM projections. Errors represent standard deviations obtained with bootstrapping.

| Bins | Eff. coverage OPES | Eff. coverage REST2 | ΔEff. coverage (%) | Norm. entropy OPES | Norm. entropy REST2 | ΔNorm. entropy (%) | Rel. coverage OPES\|REST2 | Rel. coverage REST2\|OPES | ΔRel. coverage (%) |
|---|---|---|---|---|---|---|---|---|---|
| 25 | 0.1755 ± 0.0015 | 0.1050 ± 0.0011 | -67.15 ± 1.89 | 0.7869 ± 0.0010 | 0.7211 ± 0.0011 | -9.12 ± 0.21 | 0.8626 ± 0.0095 | 0.7518 ± 0.0099 | -14.74 ± 1.84 |
| 50 | 0.1646 ± 0.0015 | 0.0979 ± 0.0011 | -68.19 ± 1.98 | 0.8150 ± 0.0008 | 0.7622 ± 0.0009 | -6.92 ± 0.16 | 0.8471 ± 0.0068 | 0.6956 ± 0.0066 | -21.77 ± 1.38 |
| 75 | 0.1555 ± 0.0013 | 0.0939 ± 0.0010 | -65.72 ± 1.92 | 0.8242 ± 0.0007 | 0.7776 ± 0.0008 | -5.99 ± 0.14 | 0.8299 ± 0.0056 | 0.6585 ± 0.0053 | -26.04 ± 1.19 |
| 101 | 0.1449 ± 0.0013 | 0.0886 ± 0.0010 | -63.61 ± 1.98 | 0.8262 ± 0.0007 | 0.7851 ± 0.0007 | -5.24 ± 0.13 | 0.8038 ± 0.0051 | 0.6302 ± 0.0045 | -27.55 ± 1.10 |
| 126 | 0.1348 ± 0.0012 | 0.0843 ± 0.0009 | -59.98 ± 1.91 | 0.8251 ± 0.0006 | 0.7879 ± 0.0007 | -4.73 ± 0.12 | 0.7750 ± 0.0047 | 0.6167 ± 0.0041 | -25.67 ± 1.03 |
| 151 | 0.1254 ± 0.0011 | 0.0789 ± 0.0009 | -58.91 ± 1.94 | 0.8223 ± 0.0006 | 0.7881 ± 0.0007 | -4.33 ± 0.11 | 0.7328 ± 0.0045 | 0.5887 ± 0.0038 | -24.47 ± 1.01 |
| 176 | 0.1154 ± 0.0010 | 0.0748 ± 0.0008 | -54.30 ± 1.86 | 0.8180 ± 0.0006 | 0.7872 ± 0.0006 | -3.91 ± 0.11 | 0.6914 ± 0.0043 | 0.5649 ± 0.0039 | -22.40 ± 1.04 |

**Table S8.** Quanitification of conformational space exploration of REST2 and OPES simulations of ACTR using the $R_g$ and helicity CVs for 2D projections. Errors represent standard deviations obtained with bootstrapping.

#### Supplementary movies

Supplementary Movies S1–S3 of the ACTR OPES simulations show trajectories 1, 2, and 5, during which multiple transitions between a disordered state and compact helical states were observed. Each movie represents 5 $\mu$s of simulation data.
